# Global changes in open reading frame dominance of RNAs during cancer initiation and progression

**DOI:** 10.1101/2023.06.02.543339

**Authors:** Yusuke Suenaga, Hiroyuki Kogashi, Kazuma Nakatani, Jason Lin, Yoshinori Hasegawa, Kazuto Kugou, Yusuke Kawashima, Eisaku Furukawa, Kazuhiro Okumura, Emiri Kita, Yuichi Wakabayashi, Mamoru Kato, Masahito Kawazu, Yoshitaka Hippo

## Abstract

Cancer cells express unique RNA transcripts; however, the factors determining their translation have remained unclear. We recently developed open reading frame (ORF) dominance as a measure that correlates with coding potential of RNAs. Upon calculating the ORF dominance of cancer-specific transcripts across 24 human tumor types, 14 exhibited significantly higher ORF dominance in cancer than in normal tissues. In organoid-based mouse genetic models, ORF dominance increased with carcinogenesis. Gene ontology analysis revealed that gene sets with increased ORF dominance were associated with cell proliferation, while those with decreased ORF dominance were linked to DNA damage response. Translatome analyses demonstrated that elevated ORF dominance during carcinogenesis resulted in higher translation frequencies of ribosome-bound RNAs. As cancer progressed, ORF dominance showed that the boundary between coding and noncoding transcripts became blurred prior to distant metastasis, indicating decreased proliferative cell populations and increased generation of RNA isoforms that potentially translate neoantigens before the development of metastatic tumors. These findings suggest that cancer evolution leads to dynamic changes in ORF dominance, resulting in global translational alterations in transcriptomes.

## Introduction

RNAs are traditionally categorized as either coding RNAs, which are translated into proteins, or noncoding RNAs. However, recent findings (Huang Y et al., 2021) have challenged this binary classification by identifying bifunctional RNAs that possess both coding and noncoding functions. For instance, our research revealed that *NCYM*, a *cis*-antisense gene of the *MYCN* oncogene, was initially classified as a noncoding RNA in the NCBI nucleotide database but is actually translated into a protein, functioning as both a coding and noncoding RNA (Suenaga et al., 2014; Shoji et al., 2015; Kaneko et al., 2015; Suenaga et al., 2020). Even a well-known algorithm (Wang et al., 2013) predicts *NCYM* to be noncoding (Suenaga et al., 2022), and an accurate method to determine its dual nature as an RNA has not yet been established. To address this issue, we developed a novel index called open reading frame dominance (ORF dominance) (Suenaga et al., 2022) that correlates with coding potential. Our investigation also revealed that while prokaryotes clearly separate the relative frequency distribution of ORF dominance between coding and noncoding RNAs, eukaryotes exhibit partial overlap, potentially facilitating the emergence of bifunctional RNAs. Moreover, we observed an increased overlap in ORF dominance distributions between coding and noncoding RNAs in endangered species, suggesting a connection to species evolution (Suenaga et al., 2022).

The relationship between the processes of carcinogenesis and biological evolution has long been debated owing to their striking similarities. Seminal studies from the 1970s demonstrated that most cancers originate from a single neoplastic cell and evolve through a selection process of somatic mutations, favoring the survival of the most proliferative population (Nowell, 1976). Subsequent research revealed that tumors consist of genetically distinct subclones with diverse characteristics, challenging the notion of linear clonal evolution (Dexter et al., 1978). The application of population genetics concepts further expanded our understanding of tumors as subjects to natural and artificial selection, including the acquisition of resistance to therapy within different subclones (Heppner, 1984). Recent advancements have introduced macroevolutionary phenomena such as chromothripsis and age-dependent carcinogenesis, which cannot be fully explained by Darwinian evolutionary theory (Vendramin et al., 2021); however, an accurate method for describing cancer evolution remains to be established.

Given that cancer tissues express RNA isoforms distinct from normal cells, which play roles in cancer initiation and progression (Vitting-Seerup et al., 2017; Kahraman et al., 2020; Karakulak et al., 2021; Climente-González et al., 2017; Kahles et al., 2018), the significance of understanding the transcriptome through RNA sequencing continues to grow. In conventional genome sequencing, short-read sequencing (SR-seq) is utilized to read shorter fragments (up to approximately 150 bp) and reconstruct the original sequence by mapping the reads to a reference. In contrast, long-read sequencing (LR-seq) has gained popularity, enabling the sequencing of tens of thousands of base pairs at once, while SR-seq is now used to refer to traditional sequencing methods. LR-seq allows direct sequencing of full-length RNA isoforms, whereas conventional SR-seq, which only maps to known references, can detect approximately 20%–40% of the total, transcript (Sharon et al., 2013; Tilgner et al., 2014). Consequently, LR-seq has facilitated the identification of novel cancer-specific isoforms across various cancer types (Chen et al., 2019; Huang KK et al., 2021; Oka et al., 2021; Fang et al., 2021; Veiga et al., 2022).

Therefore, the combination of LR-seq and ORF dominance is expected to enhance transcriptome analysis and offer new insights into cancer evolution. In light of these data, we have reevaluated cancer evolution using ORF dominance, a valuable tool for describing biological evolution.

## Materials and Methods

### ORF dominance

In this study, ORFs were defined as sequence segments that start with AUG and end with either UAA, UAG, or UGA in the 5ʹ to 3ʹ direction of the three reading frames in the RNA sequence. ORF dominance, as defined by Suenaga et al. (2022) (Figure S1), is expressed by the following formula:

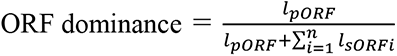

Here, *l*^pORF^ represents the length of primary ORF (pORF), and *l*_sORF_ represents the length of secondary ORF (sORF). ORF dominance values range from 0 to 1.

To calculate the ORF dominance of individual RNAs, we utilized sequence information obtained from RNA sequencing. When analyzing data from public databases or previously published articles, we determined RNA sequence information using Ensembl transcript IDs or Refseq transcript IDs and subsequently calculated ORF dominance. Ensembl transcript IDs were collected as of November 2019, while Refseq transcript IDs were collected as of April 2018. During the analysis of the LR-seq data, we identified multiple isoforms that mapped to the same Ensemble transcript ID. In such cases, we selected the longest isoform to calculate ORF dominance. Additionally, we computed the average ORF dominance of all the detected isoforms for further analysis. We plotted the relative frequency distribution of ORF dominance calculated from the entire RNA dataset and compared the peak positions and graph shapes.

### Obtaining transcriptome data

We obtained transcriptome data for ORF dominance analysis from published and public databases. Cancer-specific most dominant transcripts (cMDTs) and normal tissue transcripts (nMDTs) corresponding to cMDTs in 24 cancer types were obtained from Kahraman et al. (2020) previously published database (Table S1). To ensure high mapping accuracy, we only included data with a one-to-one correspondence, excluding cases with multiple nMDTs mapped to a cMDT. The data included mutation information for cMDTs (MutationInfo), which we used to examine the association between mutation status and ORF dominance. The mutations were classified into 6 categories based on their position: 5’UTR, CDS, 3’UTR, Core promoter, Splice site, and Enhancer. However, the data did not provide classification information for coding noncoding transcripts.

For cancer-specific transcript data obtained through LR-seq, we retrieved information on aberrant splicing of isoforms in lung adenocarcinoma specimens and lung adenocarcinoma cell lines from a previous report by Oka et al. (2021). Similarly, the classification of coding noncoding transcripts was not provided in this data. The cell line data included information on driver mutations, which we also incorporated into our analysis. Additionally, we collected data on hepatocellular carcinoma (HCC)-specific isoforms in HCC from a previous study by Chen et al. (2019), focusing on isoforms with RefSeq IDs. We divided the analysis based on coding noncoding categorization, as provided in these data. As these publications did not correspond directly to normal and cancer tissues, we used ORF dominance distributions of all coding or noncoding RNAs registered in Refseq as controls for comparison.

To examine transcript data related to neoantigens, we obtained isoform data for candidate neoantigens from a previous report by Xiang et al. (2021). Additionally, we acquired isoform data for tumor-specific antigens (TSAs) translated from noncoding gene exons from the study by Laumont et al. (2018), considering only isoforms assigned with an Ensembl transcript ID. To ensure data accuracy, we excluded isoforms with multiple Ensembl transcript IDs from the analysis.

We obtained transcriptome and clinical data of TCGA colorectal cancers from cBioPortal (https://cbioportal-datahub.s3.amazonaws.com/coadread_tcga_pub.tar.gz) and a previous report by the Cancer Genome Atlas Network (2012), respectively. The data included TNM classifications for each specimen, which we used to investigate the association between TNM classification and ORF dominance.

Transcriptome data were obtained from the study by Ono et al. (2021), which analyzed tissue diversity changes within tumors at the single-cell level using a mouse colon cancer organoid model. We separately obtained gene expression levels, sample classifications, and Ensembl transcript IDs from the authors. Samples with multiple transcripts transcribed from a single gene were excluded from the analysis since the expression level information was measured on a gene-by-gene basis.

We acquired data from a previous report by Aoto et al. (2017) on a skin carcinogenesis mice model. The gene expression levels were reported in Ensembl gene IDs, and we selected mouse genes registered in Ensembl with only one transcript ID transcribed from each gene ID for inclusion in the analysis.

### Mouse experiments

We obtained wild-type C57BL/6J mice and immunodeficient nude BALB/cA^nu/nu^ mice from Nippon Clare and Japan SLC, respectively. Kras^LSL-G12D/+^ and Trp53^flox/flox^ mice were obtained from Jackson Laboratory (Bar Harbor, ME, USA). Mice with a C57BL/6J background were bred to generate compound mutant mice with Kras^LSL-G12D/+^; Trp53^flox/flox^. Approval for safety measures to prevent proliferation and ensure animal welfare was obtained from the safety committee for recombinant DNA experiments and the animal experiment safety committee at the Chiba Cancer Center.

### Mouse organoid carcinogenesis model

We utilized a mouse organoid carcinogenesis model to recapitulate tumor formation as previously described (Ochiai et al., 2019; Matusura et al., 2020). Briefly, we established organoids from liver and pancreatic tissues from 3- to 5-week-old Kras^LSL-G12D/+^; Trp53^flox/flox^ compound mutant mice. Bile duct and pancreatic organoids were cultured in a primary culture setup. The organoids were grown in a 3D configuration between two layers of Matrigel (Corning, Corning, NY, USA) in 12- well plates. The culture medium consisted of penicillin-streptomycin (Fujifilm Wako Pure Chemical, Osaka, Japan), Fungizone (Fujifilm Wako Pure Chemical), L-glutamine (Thermo Fisher Scientific, Waltham, MA, USA), 50 ng/ml EGF (Peprotech, Rehovot, Israel), 100 ng/ml Noggin (Peprotech), 10 μM Y27632 (Fujifilm Wako Pure Chemical), and 1 μM Jagged-1 (AnaSpec, Fremont, CA, USA) in Advanced DMEM/F12 (Thermo Fisher Scientific). Passaging was performed every 4–8 days at a 1:3 dilution. We used a lentiviral vector, LV-Cre pLKO.1 (Addgene plasmid 25997), which encodes Cre-recombinase in the backbone of the pLKO.1 vector, for *in vitro* removal of the Stop codon flanked by two LoxP sequences. Successful genetic recombination was confirmed via genomic PCR. Subsequently, the organoids were subcutaneously inoculated into nude mice to induce tumor formation. The control group utilized the backbone vector pLKO.1. After approximately 8 weeks, the subcutaneous tumors were excised, and organoid culture was reestablished.

### RNA sequencing

To isolate organoid cells from the Matrigel layer, we depolymerized the Matrigel using Cell Recovery Solution (Corning). Total RNA was extracted from the isolated organoid cells using the RNeasy Mini Kit (QIAGEN, Hilden, Germany). For SR-seq, we performed sequencing on the Illumina NextSeq 500 platform and mapped the reads to the GRCm38 reference genome. LR-seq involved IsoSeq sequencing on a PacBio Sequel IIe instrument, followed by generation of Hi-Fi consensus reads. IsoSeq3 analysis was conducted using SMRTLink (version 11.0.0.146107), with the option of clustering barcoded samples separately. After IsoSeq3 analysis, the processed high-quality LR-seq sequences were mapped to the mm10 mouse genome using GMAP (version 2017-11-15). The sequences were annotated with Gencode release M25 (GRCm38.p6) using SQANTI 3 (version 5.0) with default parameters and the option of allowing the usage of transcript IDs non-related with PacBio’s nomenclature enabled. Following the process of SQANTI annotation, artifact isoforms were filtered with SQANTI3’s default rule-based filtering parameters. Major associated transcripts were sorted by their length of reference isoforms, and the longest one was selected as the representative Ensembl transcript for reporting purposes.

### Whole-genome sequencing

To isolate organoid cells from the Matrigel layer, Cell Recovery Solution (Corning) was used to depolymerize the Matrigel. DNA extraction from the isolated organoid cells was carried out using the DNeasy Blood & Tissue kit (QIAGEN). Macrogen Japan was contracted for DNA sequencing services. Whole-genome sequencing was conducted using the Illumina NovaSeq 6000 platform, and the reads were mapped to the GRCm38 reference genome.

### Western blotting

Matrigel depolymerization was performed using Cell Recovery Solution (Corning). Subsequently, organoids were lysed in RIPA (radio-immunoprecipitation assay) buffer supplemented with 1 M NaF, 0.1 M Na_3_VO_4_, 1 M βGP, and Protease Inhibitor Cocktail (Nacalai Tesque, Kyoto, Japan). The lysates were subjected to sodium dodecyl sulfate-polyacrylamide gel electrophoresis under reducing conditions, followed by transfer to polyvinylidene difluoride membranes (Millipore, Burlington, MA, USA) using a semi-dry transfer system. Membranes were blocked overnight at 4 °C in Tween buffer containing 5% dry milk for 90 min before reaction, followed by incubation with primary antibodies. The primary antibodies used were Raly (A302-070A, Bethyl Lab, 1:2,000) and α-tubulin (T5168, Sigma-Aldrich, 1:10,000).

### Creation of a list of oncogenes and tumor suppressor genes

Genes classified as oncogenes or tumor suppressor genes were identified using the UniProt database (https://www.uniprot.org/). Specifically, genes registered under the keywords "protooncogene" (KW-0656) or "tumor suppressor" (KW-0043) were categorized accordingly.

### Gene ontology analysis

We employed the DAVID Functional Annotation Tool (DAVID 2021–Dec. 2021) (https://david.ncifcrf.gov/summary.jsp) for performing gene ontology analysis (GO analysis). Gene sets were created using transcript IDs, and the Functional Annotation Chart with default settings provided by DAVID was utilized for result analysis. A Benjamini value <0.05 was considered statistically significant.

### AHA-mediated ribosome isolation

We employed the AHARIBO technique, which utilizes AHA (an analog of methionine) to incorporate it into proteins during translation. This allowed us to pull down complexes of ribosomes, RNA, and proteins involved in translation by binding them to AHA-coated beads (Minati et al., 2021). The AHARIBO kit protocol (Funakoshi, Tokyo, Japan) was followed for the experiment.

To initiate the procedure, we switched the medium of the organoids in culture to a methionine-free medium (Thermo Fisher Scientific). After 40 min of culture, AHA was added and incubated for 10 min. Following this, the medium was removed and the organoids were washed with PBS. To collect the cells from the Matrigel, the Matrigel was depolymerized using Cell Recovery Solution (Corning). The collected cells were then lysed using a kit containing lysis buffer. The ligand-coated beads were bound to the lysate, and the resulting bead-cell complex was collected using a magnetic rack. The collected bead suspension was incubated with 10% Sodium Dodecyl Sulfate and Proteinase K (Fujifilm Wako Pure Chemical) at 37 °C for 60 min. The supernatant was collected, and RNA was extracted using RNeasy mini (QIAGEN) for RNA sequencing. Similarly, for protein analysis, the beads were reacted with the cell lysate and collected using a magnetic rack. The collected bead suspension was subjected to LC-MS analysis following a thorough wash with urea washing solution (AHARIBO kit).

### Quantitative real time RT-PCR

RNA was extracted using the AHARIBO kit, and cDNA synthesis was performed using SuperScript II with random primers (Invitrogen, Waltham, MA, USA). Quantitative real-time PCR was conducted on a StepOnePlus™ Real-Time PCR System (Thermo Fisher Scientific) with SYBR green. The following primer sets were used: *Raly* (5′-GTGCTCGGGCTCCTCAC-3′ and 5′-GGACATGGTGTTCACCCGC-3′) and *Rn18s* (5′-GTAACCCGTTGAACCCCATT-3′ and 5′-CCATCCAATCGGTAGTAGCG-3′). The expression of the *Raly* gene was normalized to *Rn18s* RNA levels.

### LC-MS

To digest proteins, we added 500 ng of trypsin/Lys-C Mix (CAT# V5072, Promega, Madison, WI, USA) to the beads soaked in 50 mM Tris-HCl pH 8.0. The mixture was gently mixed at 37 °C overnight. The resulting digested sample (supernatant) was collected in a new 1.5 ml tube. The collected sample was then treated with 20 mM tris (2-carboxyethyl) phosphine at 80 °C for 10 min for reduction, followed by alkylation using 30 mM iodoacetamide at room temperature for 30 min while protecting it from light. The alkylated sample was acidified with 20 μL of 5% trifluoroacetic acid (TFA) and desalted using a STAGE tip (CAT# 7820-11200, GL Sciences, Tokyo, Japan) according to the manufacturer’s protocol. After desalting, the sample was dried using a centrifugal evaporator (miVac Duo concentrator, Genevac, Ipswich, UK). The dried sample was then dissolved in 2% ACN containing 0.1% TFA.

For peptide analysis, the peptides were directly injected onto a 75 μm × 12 cm nanoLC nano-capillary column (Nikkyo Technos, Tokyo, Japan) at 50 °C. Separation was achieved using a 60 min gradient at a flow rate of 200 nl/min with an UltiMate 3000 RSLCnano LC system (Thermo Fisher Scientific). Peptides eluting from the column were analyzed using a Q Exactive HF-X (Thermo Fisher Scientific) for DIA-MS. MS1 spectra were collected in the range of 495–745 m/z at a resolution of 15,000, with an automatic gain control (AGC) target of 3e6 and a maximum injection time of 23 ms. MS2 spectra were collected at a resolution of 30,000 in the range of more than 200 m/z, with an AGC target of 3e6, maximum injection time set to "auto," and a normalized collision energy of 23%. The isolation width for MS2 was set to 4 m/z, and window placements optimized by Scaffold DIA 3.2.1 were used with window patterns of 500–740 m/z.

For MS file analysis, we used DIA-NN (version: 1.8.1, https://github.com/vdemichev/DiaNN) (Demichev, et al., 2020) to search against the in silico mouse spectral library. The spectral library was generated from the mouse protein sequence database (proteome ID UP000000589, 21,986 entries, downloaded on November 26, 2021) using DIA-NN. Parameters for generating the spectral library were as follows: precursor m/z range of 490–750; fragment ion m/z range of 200– 1,800; precursor charge range of 2–4; peptide length range of 7–45; digestion enzyme of trypsin; missed cleavages of 1; and enabled features included "C Carbamidomethylation," "n-term M excision," "deep learning-based spectra, RTs and IMs prediction," and "FASTA digest for library-free search/library generation." The DIA-NN search parameters included neural network classification in single-pass mode, quantification strategy of robust LC (high precision), mass accuracy of 10 ppm, MS1 accuracy of 10 ppm, protein inference based on genes, and enabled features such as "MBR," "no shared spectra," "unrelated runs," "use isotopologues," and "heuristic protein inference." The protein identification threshold was set at 1% or less for both precursor and protein FDRs.

## Results

### Changes in ORF dominance during carcinogenesis

To gain insights into the significance of changes in ORF dominance during carcinogenesis, we calculated the ORF dominance of transcriptomes obtained from normal and cancer tissues and compared their distributions.

Using a dataset of 24 cancer types from Kahraman et al. (2020), which included cancer-specific cMDTs and corresponding nMDTs in normal tissues, we initially compared the ORF dominance distributions of all cMDTs and nMDTs regardless of cancer type. We observed a significant shift toward higher ORF dominance values in cMDTs compared to nMDTs (Figure 1A). Next, we focused on transcripts derived from oncogenes and tumor suppressor genes within cMDTs and compared them to nMDTs. We found that the ORF dominance of cMDTs significantly increased only for oncogenes (Figure 1B). Additionally, when comparing ORF dominance by cancer type, cMDTs showed significantly higher values in 14 out of 24 cancer types, while three cancer types (bladder transitional cell carcinoma (Bladder-TCC), chronic lymphocytic leukemia (Lymph-CLI), bone and soft tissue leiomyosarcoma (Bone-Leiomyo)) exhibited lower ORF dominance. No significant differences were detected among seven other cancer types (Figure 1C).

**Figure 1.**
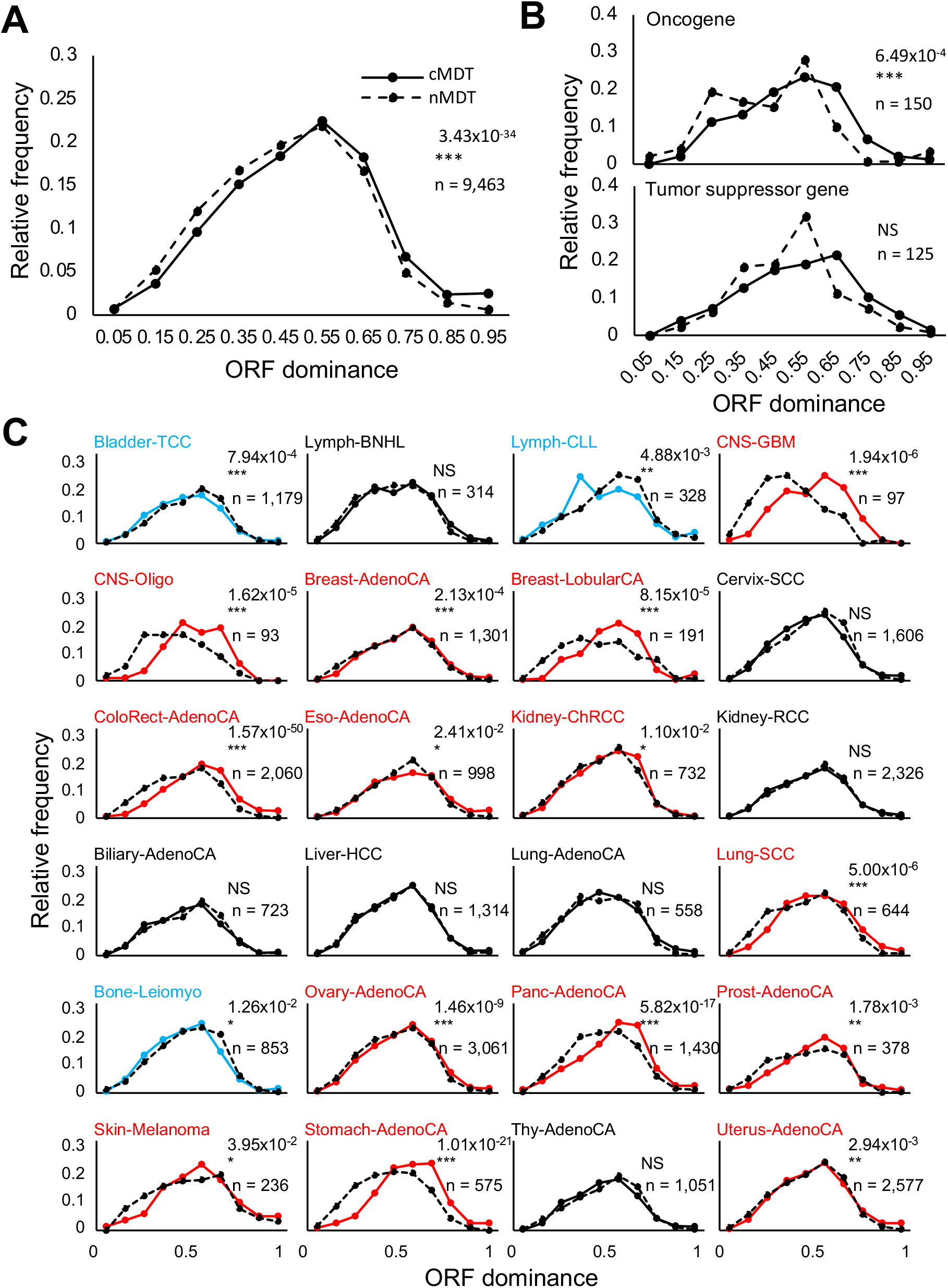
Distribution of ORF dominance in cancer-specific transcripts across 24 cancer types. Comparison between cMDT and nMDT. (A) Distribution of ORF dominance in all transcripts expressed in 24 cancer types. Solid line represents cMDT, dashed line represents nMDT. (B) Distribution of ORF dominance in oncogenes and tumor suppressor genes. (C) Changes in ORF dominance in cMDT indicated by increased (red line) or decreased (blue line) values. *P*-value was determined using the Mann–Whitney *U* test (***: *P* < 0.001, **: *P* < 0.01, *: *P* < 0.05, NS: not significant).

These findings indicate an overall increasing trend in transcript ORF dominance during carcinogenesis, specifically elevated ORF dominance for oncogenes and cancer type-specific transcripts.

Furthermore, GO analysis of gene sets with ORF dominance changes exceeding 0.1 between nMDT and cMDT revealed that genes with increased ORF dominance were associated with the Golgi apparatus, endoplasmic reticulum, cell junctions, and cell cycle (Table S2), while those with decreased ORF dominance were related to the mitochondrion, DNA damage, and DNA repair (Table S3). Similarly, analyzing the 14 cancer types with increased ORF dominance showed that gene sets with elevated ORF dominance were associated with the Golgi apparatus, endoplasmic reticulum, and cell junctions, whereas those with decreased ORF dominance were linked to the mitochondrion, DNA damage, and DNA repair (Table S4, S5). Likewise, for the three cancer types with decreased ORF dominance (Bladder-TCC, Lymph-CLI, and Bone-Leiomyo), the gene sets with decreased ORF dominance were associated with the mitochondrion (Table S6). These results suggest that changes in ORF dominance may be associated with physiological functions involved in carcinogenesis, such as organelles, the cell cycle, and DNA damage response.

To examine the relationship between changes in ORF dominance and genetic mutations, we compared the ORF dominance distributions of nMDT and cMDT among isoforms with and without mutations. The results showed a significant shift toward higher ORF dominance values irrespective of the presence or absence of mutations (Figure S2A). Additionally, investigating the association between the change in ORF dominance and mutation location revealed a significant increase in ORF dominance when mutations were present in the 5’ UTR, 3’ UTR, and CDS regions. However, no change in ORF dominance was detected when mutations occurred in regulatory regions such as the core promoter, enhancer, and splice sites (Figure S2B).

### LR-seq reveals high-resolution ORF dominance changes

The data presented above are derived from SR-seq data obtained from the Pan-Cancer Analysis of Whole Genomes (PCAWG) project (Kahraman et al., 2020). Notably, no detectable changes in ORF dominance were observed in lung adenocarcinoma (Lung-AdenoCA) and hepatocellular carcinoma (Liver-HCC) based on these findings. However, recent studies utilizing LR-seq (Oka et al., 2021; Chen et al., 2019) have identified cancer-specific isoforms in samples and cell lines of these cancer types. Consequently, we reevaluated ORF dominance using these new datasets and observed a significant shift toward higher values in numerous lung adenocarcinoma samples and cell lines (Figure S3). In the lung adenocarcinoma cell line data, all cell lines carrying *KRAS* mutations exhibited a marked shift in ORF dominance, whereas those with *NRAS* or *EGFR* mutations did not demonstrate a significant change (Figure S3A). Conversely, in hepatocellular carcinoma, a significant shift toward lower values in ORF dominance was observed (Figure S3B). These outcomes indicate that LR-seq exhibits high sensitivity in detecting changes in ORF dominance.

Moreover, recent studies (Xiang et al., 2021; Laumont et al., 2018) have identified neoantigens translated from cancer-specific isoforms across different cancer types. Since ORF dominance was originally developed as an indicator of coding potential encompassing noncoding transcripts (Suenaga et al., 2022), it is likely to serve as a valuable predictor of neoantigens. To validate this, we calculated the ORF dominance based on the data reported in the aforementioned studies and observed significantly high ORF dominance in the isoforms of each neoantigen candidate (Figure S4).

### Changes in ORF dominance in a mouse model of organoid carcinogenesis

To examine the changes in ORF dominance during pure carcinogenesis, we compared normal and cancer cells with matched backgrounds derived from the same mouse cells. Specifically, we lentivirally transduced *Kras*^LSL-G12D/+^; *Trp53*^flox/flox^ mouse bile duct organoids *in vitro* with either Cre or the empty vector pLKO.1 to obtain *Kras*^G12D/+^; *Trp53*^-/-^ (hereafter referred to as Cre organoids) and normal controls (V organoids). When both types of organoids were inoculated into nude mice, subcutaneous (SC) tumors only developed from the Cre organoids. We obtained two clones of subcutaneous tumor-derived organoids (SC1/SC2 organoids) from two independent experiments (Figure 2A-C). Total RNA was extracted from these four organoids, and RNA sequencing (LR-seq) was performed. In the analysis, transcripts were classified as coding RNA or noncoding RNA during annotation, and the distribution of ORF dominance was analyzed for each.

**Figure 2.**
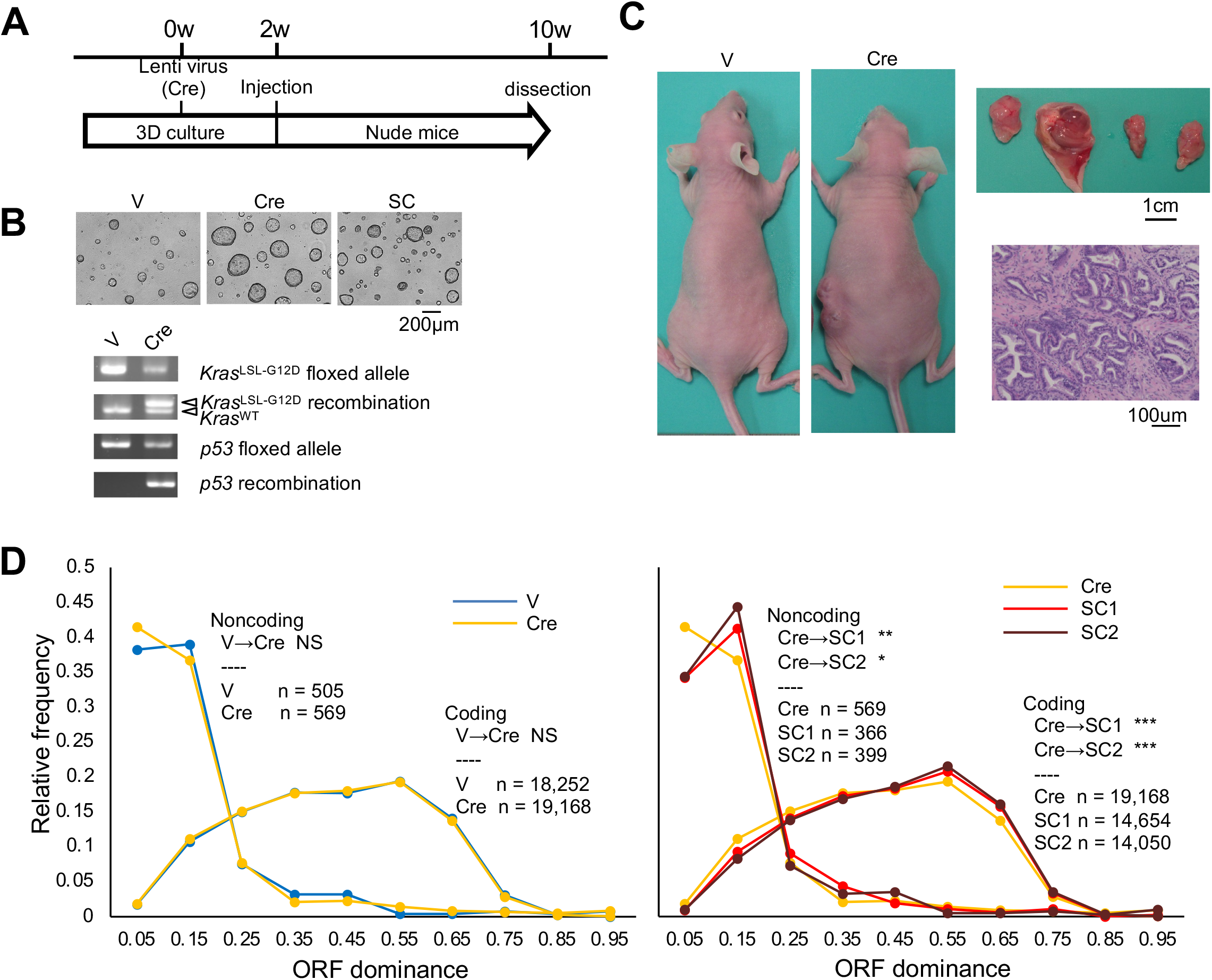
Shift in ORF dominance distributions during carcinogenesis. (A) Experimental protocols for genetic manipulation of mouse organoids. (B) Phase contrast images of mouse organoids and genomic PCR analysis focused on *Kras* and *p53* loci. (C) Representative images of developed tumors and hematoxylin & eosin (H&E) staining. V: vehicle, Cre: mutation-induced organoid by Cre recombinase. SC1/2: Subcutaneous tumor-derived organoid. (D) ORF dominance distribution of transcripts based on LR-seq data. *P*-value was determined using the Mann–Whitney *U* test (***: *P* < 0.001, **: *P* < 0.01, *: *P* < 0.05, NS: not significant).

The results showed no significant differences in the ORF dominance distribution of the organoids before (V) and after (Cre) genetic engineering, for both coding and noncoding RNAs. However, the ORF dominance distribution of the transcripts before (Cre) and after tumorigenesis (SC1, SC2) exhibited a significant shift toward higher values for both coding and noncoding RNAs (Figure 2D and Figure S5). Additionally, GO analysis of gene sets with ORF dominance changes greater than 0.1 revealed that gene sets with increased ORF dominance were associated with Golgi apparatus, endoplasmic reticulum, mitosis, cell division, and cell cycle (Table S7), similar to the results obtained in human cancers. Gene sets related to mRNA processing, DNA damage, and DNA repair were enriched in gene sets with both increased and decreased ORF dominance (Table S7 and S8).

Additionally, we performed similar analysis using mouse pancreatic organoids with the *Kras*^LSL-G12D/+^; *Trp53*^flox/flox^ allele to obtain V, Cre, and SC organoids as previously described in our study (Matsuura et al., 2020). After LR-seq analyses, we examined the ORF dominance distributions. The ORF dominance distributions after genetic engineering (Cre) were significantly shifted toward higher values in both coding RNA and noncoding RNA, compared to the distributions in V organoids (Figure S6). Similar to human cancers and cholangiocarcinoma organoid models, the gene sets with increased ORF dominance after genetic engineering (Cre) were associated with Golgi apparatus, endoplasmic reticulum, cell junction, and cell cycle (Table S9). Gene sets related to mRNA processing, mitochondrion, and autophagosomes were enriched in gene sets with both increased and decreased ORF dominance (Table S9 and S10). After tumorigenesis (SC), gene sets related to Golgi apparatus and endoplasmic reticulum were enriched in genes with both increased and decreased ORF dominance (Table S11 and S12), while gene sets related to cell cycle, cell division, and mitosis were predominantly enriched in decreased ORF dominance. These results indicate that the timing of ORF dominance elevation differs among different tissues even when the same genetic mutations are employed for carcinogenesis. Moreover, certain gene sets with altered ORF dominance share common functions (e.g., Golgi apparatus, endoplasmic reticulum, and cell cycle), while others are tissue-specific (e.g., autophagosomes).

### RNA sequence changes inducing alternations of ORF dominance

Using the LR-seq method in a mouse bile duct organoid model, we investigated changes in ORF dominance for all transcripts during the process of carcinogenesis. The analysis revealed that the percentage of transcripts with increased ORF dominance rose during *in vivo* tumorigenesis (Cre→SC) compared to *in vitro* genetic engineering (V→Cre), while the percentage of transcripts with decreased ORF dominance declined during tumorigenesis (Cre→SC) compared to genetic engineering (V→Cre) (Figure S7A). Approximately 50% of all transcripts exhibited altered ORF dominance during tumorigenesis (Cre→SC), whereas around 40% of the transcripts showed altered ORF dominance at the time of genetic engineering (V→Cre), with a similar rate of increase and decrease. Consequently, the overall distribution of ORF dominance remained unchanged during genetic engineering (V→Cre) (Figure 2D). Notably, a significant proportion of transcripts with elevated ORF dominance during tumorigenesis (Cre→SC) resulted from a reduction in the cumulative length of all sORFs, rather than elongation of the pORF (Figure S7B, left). Conversely, transcripts with decreased ORF dominance were more likely to be attributed to pORF shortening (Figure S7B, right).

It is known that shortening of the 3’ untranslated region (3’UTR) enhances the expression of proteins encoded by oncogenes (Mayr et al., 2009), while intronic polyadenylation contributes to the inactivation of tumor suppressor genes and DNA repair genes (Lee et al., 2018; Dubbury et al., 2018). Therefore, we further analyzed the causes of increased and decreased ORF dominance by focusing on oncogenes and tumor suppressor genes, but the results did not significantly differ from the overall analysis (Figure S7C). Enrichment analysis of gene sets with increased ORF dominance and shortened cumulative sORF lengths, as well as gene sets with decreased ORF dominance and shortened pORF, revealed an enrichment of genes related to the Golgi apparatus, endoplasmic reticulum, mitosis, cell cycle, cell division, DNA repair, and DNA damage (Table S13 and S14).

To explore the potential contributions of genomic mutations to the elevation of ORF dominance after tumorigenesis, whole-genome sequencing was performed on the four organoids (Figure S8). The number of mutations increased following tumorigenesis (Figure S8A); however, no significant changes were observed in the types of mutations (Figure S8B).

### Elevation of ORF dominance contributes to enhanced translation

In our previous study (Suenaga et al., 2022), we reported an association between ORF dominance and translation efficiency during spermatogenesis in the testis. Based on this finding, we hypothesized that an increase in ORF dominance may lead to enhanced translation during carcinogenesis, potentially resulting in increased protein expression. To investigate this, we examined genes exhibiting a significant increase in ORF dominance during tumorigenesis (Cre→SC) in a mouse cholangiocarcinoma model (Table S15). Among these genes, we selected *Raly* due to its known oncogenic functions (Sun et al., 2020; Cornella et al., 2017) and its mRNA levels showing no significant changes (Table S15 and Figure 3A). After tumorigenesis (SCs), the ORF dominances of *Raly* mRNA increased due to 3’UTR shortening (Figure 3A), and this was accompanied by elevated protein expression levels (Figure 3B). To further investigate the translation process, we extracted RNAs bound to ribosomes, which were actively translating nascent peptides, from the organoids using AHARIBO. Subsequently, we quantified the *Raly* mRNA bound to ribosomes and the newly translated Raly protein using RT-qPCR and LC-MS, respectively. While the amount of *Raly* mRNA bound to ribosomes in SCs showed a decrease compared to Cre (Figure 3C), the level of newly translated Raly protein was higher in SCs (Figure 3D), indicating that increased ORF dominance is associated with augmented translation of Raly.

**Figure 3.**
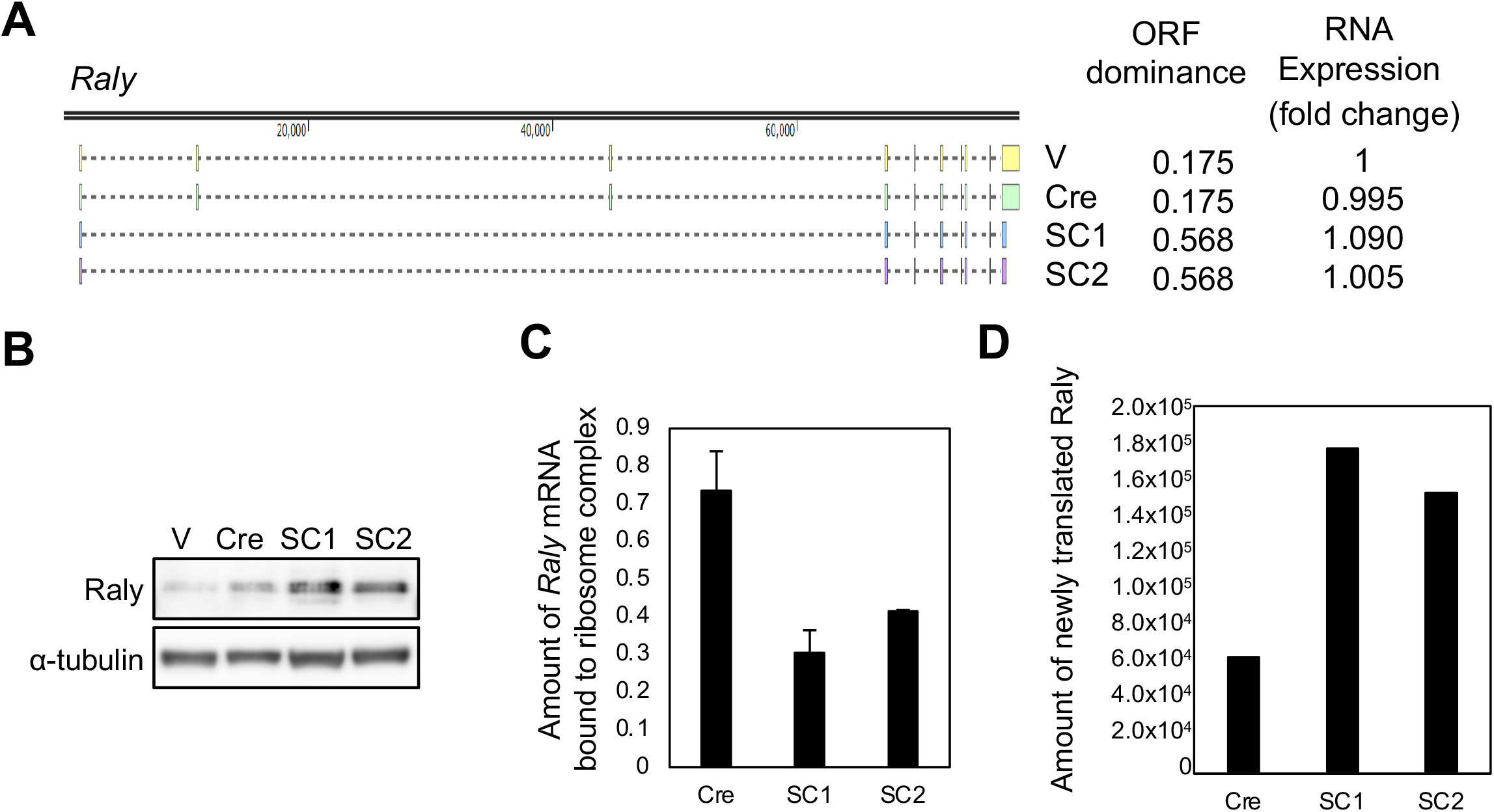
Enhanced translation of Raly protein due to increased ORF dominance of *Raly* transcripts. (A) Structure of *Raly* gene (left). Isoforms expressed in indicated organoids, along with their corresponding ORF dominance and mRNA expression levels (right). Dot lines and colored rectangles represent introns and exons, respectively. mRNA expression levels were assessed by SR-seq, and fold changes were calculated relative to the expression levels in V. (B) Western blotting showing expression levels of Raly protein in the indicated organoids. α-tubulin was used as a loading control. (C) Amount of *Raly* mRNA bound to ribosome complexes measured by RT-qPCR after AHARIBO. Values were normalized to ribosome RNA, *Rn18s*. (D) Amount of newly translated Raly protein analyzed by LC-MS after AHARIBO. Total protein amounts were normalized before AHARIBO, and relative protein levels were determined by LC-MS.

To gain a comprehensive understanding of the relationship between ORF dominance and translation, we conducted a thorough analysis of RNAs bound to ribosomes that were actively translating nascent peptides using AHARIBO combined with LR-seq. We also calculated the ORF dominance based on the RNA sequences (Figure 4A and Figure S9). The distributions of ORF dominance for ribosome-bound RNAs revealed a shift toward lower values in coding transcripts (Figure 4A, left), while noncoding transcripts showed a shift toward higher values, although the shift in SC2 noncoding RNAs was not statistically significant (Figure 4A, right). The elevation of ORF dominance following tumorigenesis in coding transcripts (Cre < SCs, bulk) was mirrored in RNAs bound to translating ribosomes (Cre < SC1 < SC2, AHA). However, the elevation of noncoding RNAs (Cre < SCs, bulk) was only evident in SC1 noncoding RNAs bound to ribosomes, not in SC2 (Cre ≈ SC2 < SC1, AHA). These results suggest that changes in ORF dominance within the transcriptome are only partially reflected in RNAs bound to translating ribosomes, and the extent of this reflection varies across different tumors. The prominent shift in coding RNAs was observed in SC2, while the shift in noncoding RNAs was notable in SC1.

**Figure 4.**
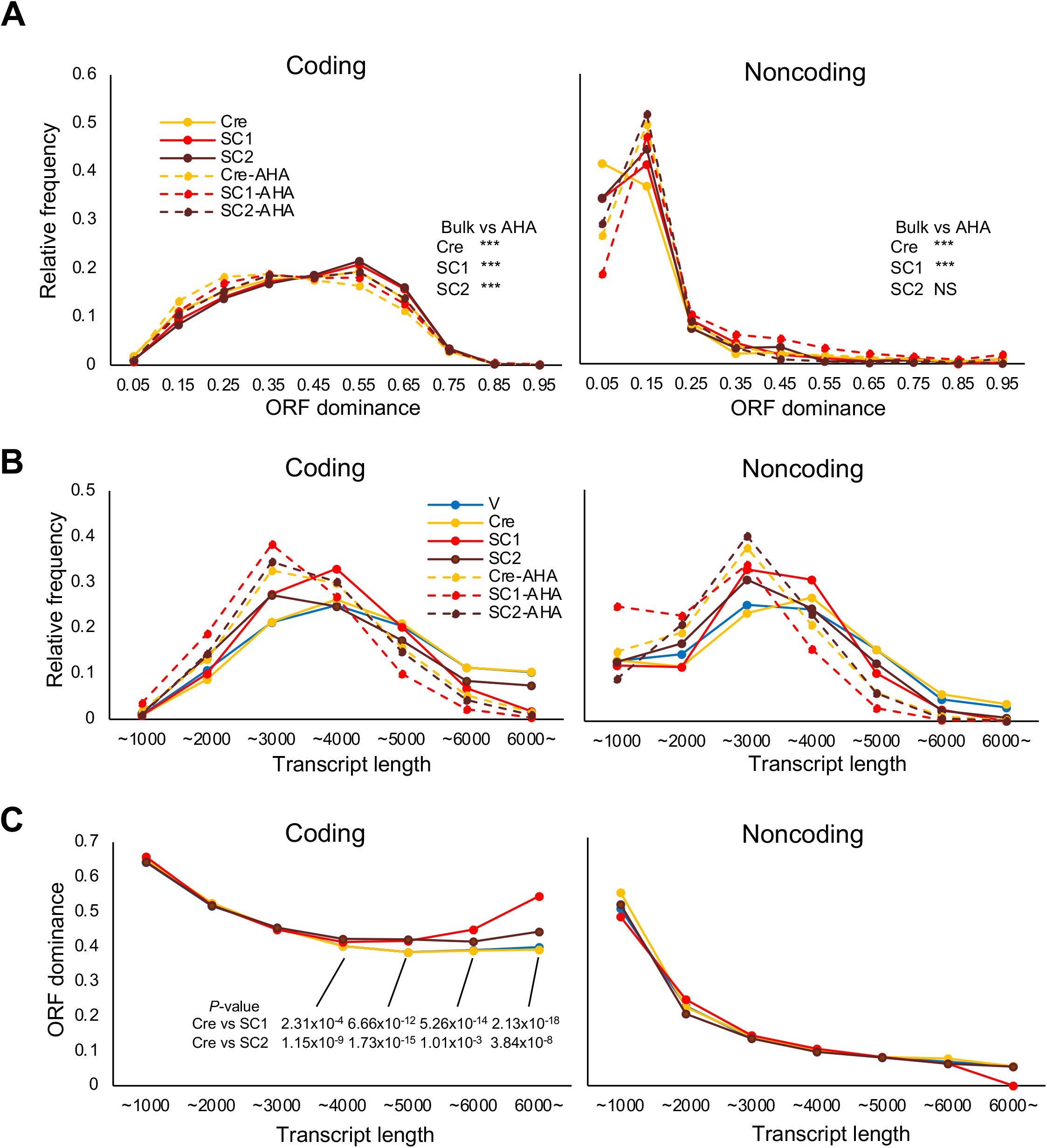
Shift in ORF dominance of RNAs bound to ribosome complexes in mouse cholangiocarcinoma organoids. (A) ORF dominance distributions of coding (left) and noncoding (right) transcripts bound to ribosome complexes (AHA) in mouse bile duct organoids. Bold and dotted lines indicate all RNAs detected in the organoids (bulk, same data as Figure 2D) and RNAs detected after AHARIBO (AHA), respectively. Statistical significance was determined using the Mann–Whitney *U* test (***: *P* < 0.001, NS: not significant). (B) Relative frequencies of transcript length (nucleotide) in coding (left) and noncoding (right) transcripts. (C) Relationship between transcript length and ORF dominance in coding (left) and noncoding (right) transcripts. *P*-value was determined using Student’s *t*-test.

To investigate the mechanism underlying the shifts in ORF dominance upon ribosome binding, we examined other RNA features, such as transcript length and expression levels, as they are related to ribosome binding. Shorter transcripts have been associated with higher ORF dominance (Suenaga et al., 2022), whereas longer transcripts are linked to increased ribosome binding (Zeng and Hamada, 2018). This suggests the presence of an optimal transcript length for efficient translation. Consistently, the relative frequencies of transcript length for RNAs bound to translating ribosomes showed peaks around 3,000 bases for both coding and noncoding transcripts (Figure 4B, AHA). Additionally, cancer organoids (SCs) exhibited transcriptomes that approached this optimal transcript length compared to control samples (V and Cre). As previously reported, shorter transcripts demonstrated higher ORF dominance in both coding and noncoding transcripts (Figure 4C). Shortening transcripts was the primary mechanism for achieving optimal translation (Figure 4B). However, in cancer organoids, long coding transcripts (>4,000 bp) showed increased ORF dominance, whereas noncoding transcripts did not exhibit the same pattern (Figure 4C). This difference between coding and noncoding transcripts is likely due to the presence of functional ORFs in coding transcripts, which limits transcript shortening. No relationship was observed between RNA expression levels and ORF dominance or tumorigenesis (Figure S10 and S11).

Ribosomes translate peptides from both pORFs and sORFs. Our methodology (AHARIBO-LR-seq) allowed us to extract and identify all RNAs translated from pORFs and/or sORFs. However, AHARIBO-LC-MS data specifically identified peptides from *bona fide* ORFs, as peptides from sORFs were either not registered in the database or were undetectable due to low stability. Consequently, we examined whether ORF dominance-shifts in ribosome-bound RNAs contributed to the translation of proteins from *bona fide* ORFs (Figure 5). The number of transcripts with evidence of translation from *bona fide* ORFs (+) was substantially higher than in those without evidence (-) in cancer organoids (SCs) (Figure 5A), aligning with the order of ORF dominance-shifts in RNAs bound to translating ribosomes (Figure 4A, Cre < SC1 < SC2, AHA). The reduction in transcripts without evidence of translation (-) in cancer organoids (Figure 5A) also corresponds to the observed decrease in sORFs (Figure S7B), indicating that the primary mechanism driving elevated ORF dominance after tumorigenesis involves a reduction in sORFs. Furthermore, the relationship between translational frequencies (the frequency of translation from *bona fide* ORFs) and ORF dominance approximated a linear function passing through 0.3 when ORF dominance was below 0.7 in cancer organoids (SCs) (Figure 5B), similar to the linear relationship observed between ORF dominance and coding potential within the same range (<0.7) (Suenaga et al., 2022). However, the linear relationship was not detected in the control (Cre) and the basal levels of translational frequencies were approximately 20% lower than those of cancer organoids. Only transcripts with the highest ORF dominance (> 0.9) in Cre showed comparable translational frequencies to those in SCs. Therefore, during carcinogenesis, the basal level of translation from RNA bound to ribosomes increased and ORF dominance-dependency of translational frequencies emerged. The amount of newly translated proteins per gene was higher in cancer organoids (Figure 5C), reflecting the order of ORF dominance shifts (Figure 4A) and translational frequencies (Figure 5A) (Cre < SC1 < SC2). Notably, this elevation was only observed within the defined range of ORF dominance (< 0.7) (Figure 5B). These findings indicate that shifts in ORF dominance within the transcriptome contribute to enhanced translation in cancer organoids.

**Figure 5.**
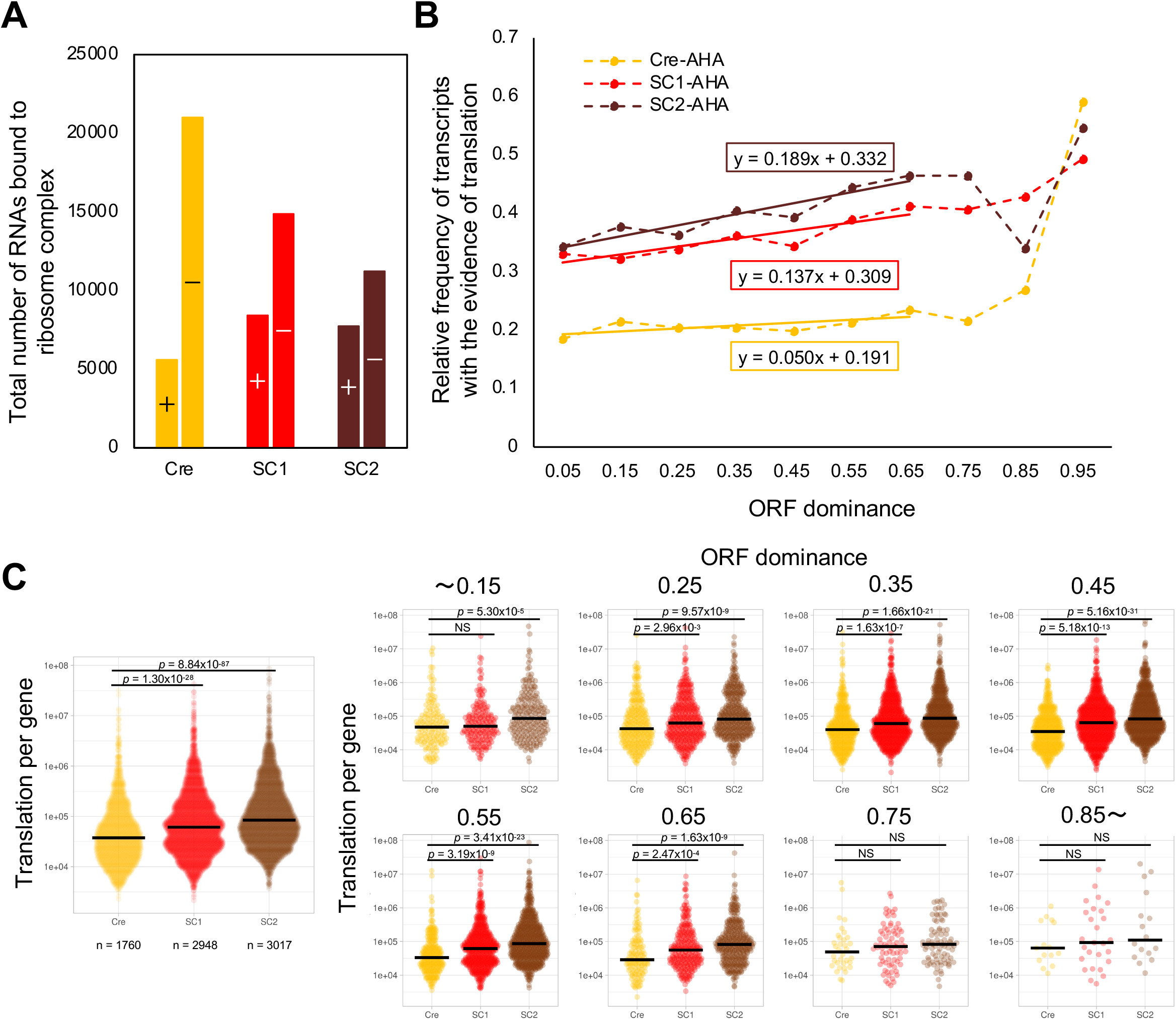
Relationship between ORF dominance and translation in the mouse cholangiocarcinoma model. (A) Total numbers of coding transcripts with (+) or without (-) evidence of translation from a bona fide ORF in mouse bile duct organoids. *P*-value was determined using Fisher’s exact test. (B) Linear relationship between ORF dominance and relative frequency of coding transcripts with evidence of translation from a bona fide ORF. *P*-value was determined using Fisher’s exact test (***: *P* < 0.001, *: *P* < 0.05, NS: not significant). (C) Translation level per gene in the indicated organoids. Statistical significance was determined using the Mann–Whitney *U* test. Violin plots were generated using PlotsOfData (Postma *et al*., 2019).

### Potential of ORF dominance analysis using SR-seq data

To validate the efficacy of LR-seq in ORF dominance analysis, we compared the ORF dominance distributions using SR-seq data from the same RNA samples in a mouse cholangiocarcinoma model experiment (Figure S12). The SR-seq results revealed no significant difference in ORF dominance distribution between the four organoids for both coding and noncoding RNAs (Figure S12A). This led us to consider the possibility that RNAs with unchanged expression levels might introduce noise into the analysis, masking the results of RNAs with altered ORF dominance. To address this, we extracted RNAs with expression levels increased more than two-fold before and after genetic engineering (V→Cre) and tumorigenesis (Cre→SC1, Cre→SC2). We then compared the ORF dominance distributions again. The analysis of noncoding RNAs showed no significant difference, whereas in the analysis of coding RNAs, the ORF dominance distribution of RNAs with expression level changes after tumorigenesis (Cre→SC1, Cre→SC2) was higher than that of RNAs with expression level changes after genetic engineering (V→Cre) for upregulated RNAs (Figure S12B). These findings suggest that LR-seq can more accurately detect changes in ORF dominance compared to SR-seq. Furthermore, our results indicate that even with SR-seq data, changes in ORF dominance can be detected with high sensitivity by focusing on transcript data with altered RNA expression levels.

To explore the relationship between ORF dominance and intratumor diversity, we calculated ORF dominance using single-cell RNA sequencing data obtained via SR-seq (Ono et al., 2021). In a previous study, mouse colorectal cancer organoids were derived from single cells with APC knockdown and subcutaneously inoculated into nude mice to induce tumor formation. The organoids were then cultured again from the tumors and subjected to subcutaneous inoculation for secondary tumor formation and organoid reculturing. The organoid before inoculation into nude mice was referred to as time point 1 (Tp1), the organoid derived from the first tumor as Tp2, and the organoid derived from the second tumor as Tp3. Cell populations from each organoid were classified into three groups based on the expression levels of a group of marker genes: AntiEpi, cGMP/GC, and Dormant (Ono et al., 2021). AntiEpi represents an ancestral type persisting throughout cancer evolution (Tp1 to Tp3), while cGMP/GC and Dormant-types are newly emerged populations after Tp1. Human colorectal cancers with expression patterns resembling cGMP/GC and Dormant were associated with distant metastasis (Ono et al., 2021). We examined the expression levels of the three types at Tp2 or Tp3 compared to AntiEpi at Tp1 and compared the ORF dominance distribution of transcripts with more than a two-fold increase in expression levels to those with no expression change (Figure 6A). No significant difference was observed at the bulk level (Figure 6A, top), but elevated ORF dominance was observed in the newly emerged populations at Tp2 (Figure 6A, left). Specifically, cGMP/GC showed elevated ORF dominance in both coding and noncoding transcripts, while Dormant-type showed higher ORF dominance only in coding transcripts. At Tp3, even AntiEpi showed higher ORF dominance in both coding and noncoding transcripts compared to Tp1 (Figure 6A, right). These results suggest that an increase in ORF dominance occurs in newly emerged cell populations (cGMP/GC and Dormant) followed by an increase in ORF dominance in pre-existing cell populations (AntiEpi). Human colorectal cancers were classified into three subgroups (Ono et al., 2021), and we calculated the ORF dominance of transcripts with increased expression in the cGMP/GC or Dormant subgroups compared to the AntiEpi subgroup (Figure 6B). ORF dominance of coding transcripts was elevated in the cGMP/GC or Dormant subgroups, while no significant differences were observed in noncoding transcripts. Therefore, consistent with the mouse model, an elevation in ORF dominance was observed in the metastatic subgroups (cGMP/GC and Dormant) of human colorectal cancers.

**Figure 6.**
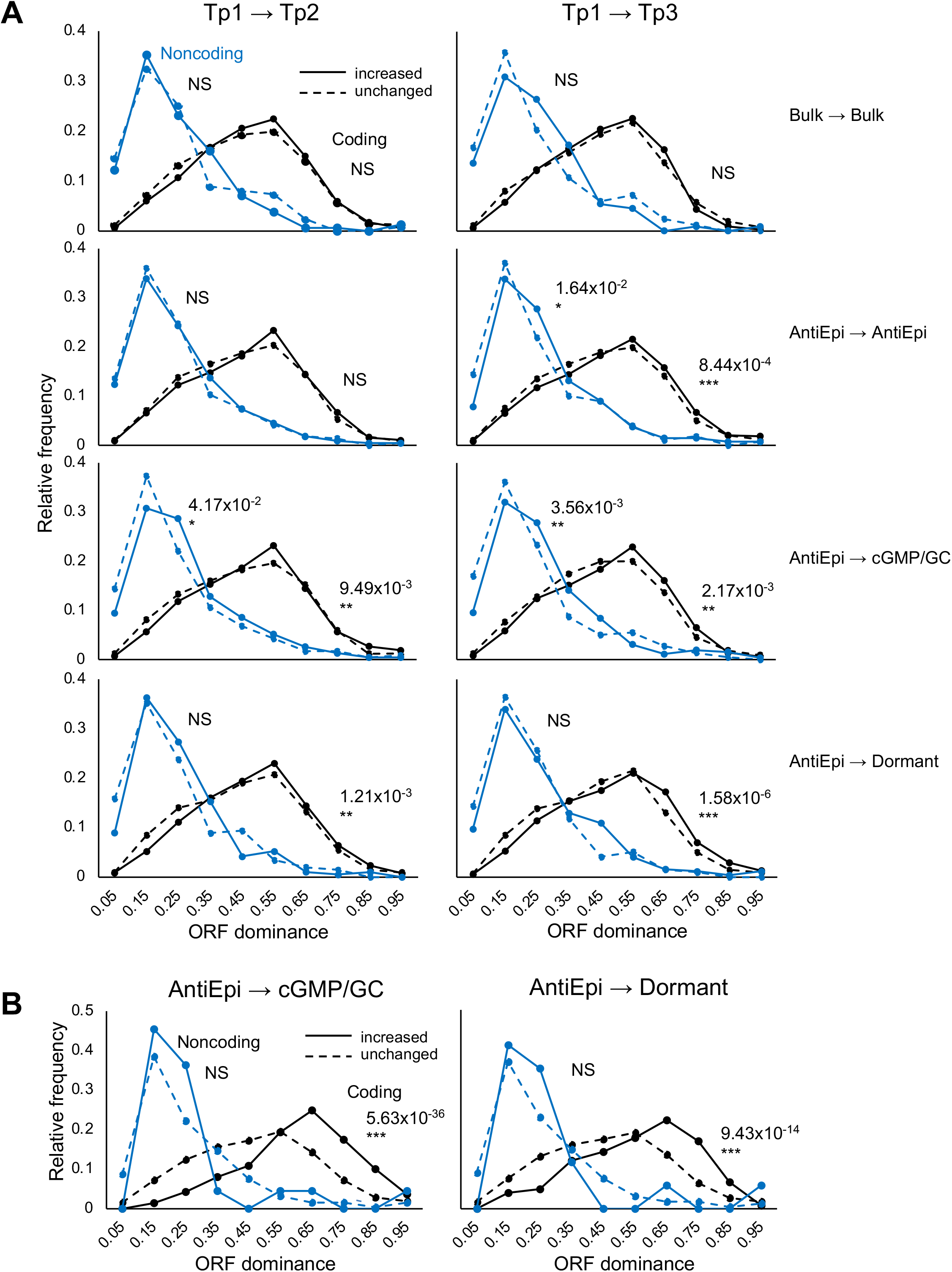
ORF dominance distribution in three subtypes of mouse and human colorectal cancers. (A) ORF dominance distributions of transcripts in mouse colorectal carcinoma model with increased (solid line) or unchanged (dashed line) expression in three subtypes at time points 2 (left) or 3 (right), compared to Anti-epithelial subtype (AntiEpi) at time point 1. Bulk tissue data at time points 2 (top left) and 3 (top right) were compared to bulk data at time point 1. Tp: time point. (B) ORF dominance distribution of transcripts with increased (solid line) or unchanged expression in cGMP/GC (left) or Dormant (right) subgroups, compared to AntiEpi subgroup in human colorectal cancers. *P*-value was determined using the Mann–Whitney *U* test (***: *P* < 0.001, **: *P* < 0.01, *: *P* < 0.05, NS: not significant).

To investigate the relationship between ORF dominance and cancer progression, we calculated ORF dominance based on TNM classification (Table S16). We determined the ORF dominance of transcripts with increased expression levels in late stages (T1,2 vs T3,4; N0 vs N1, 2; M0 vs M1) within the three subgroups (Figure 7). Similar to the mouse model (Figure 6A), a prominent elevation in ORF dominance of noncoding transcripts was observed only in the cGMP/GC subtype at late stages of T and N. In contrast, the peaks of the ORF dominance distribution of coding transcripts shifted to lower values in AntiEpi and cGMP/GC at the late stages of T and N (Figure 7). Notably, no significant change in ORF dominance with cancer progression was detected in the Dormant subtype, which has the highest frequency of distant metastases (Ono et al., 2021). The ORF dominance distributions of coding transcripts remained unchanged in AntiEpi and cGMP/GC regardless of the presence of metastatic cancers. Transcript length remained largely unchanged during cancer progression, except for shorter noncoding and longer coding transcripts in the cGMP/GC subgroup at advanced stages of T and M, respectively (Figure S13). Based on these findings, we hypothesized that during cancer progression, there is an increase in ORF dominance in noncoding transcripts and a decrease in coding transcripts. These shifts in ORF dominance occur before the emergence of metastatic cancers without changes in transcript length.

**Figure 7.**
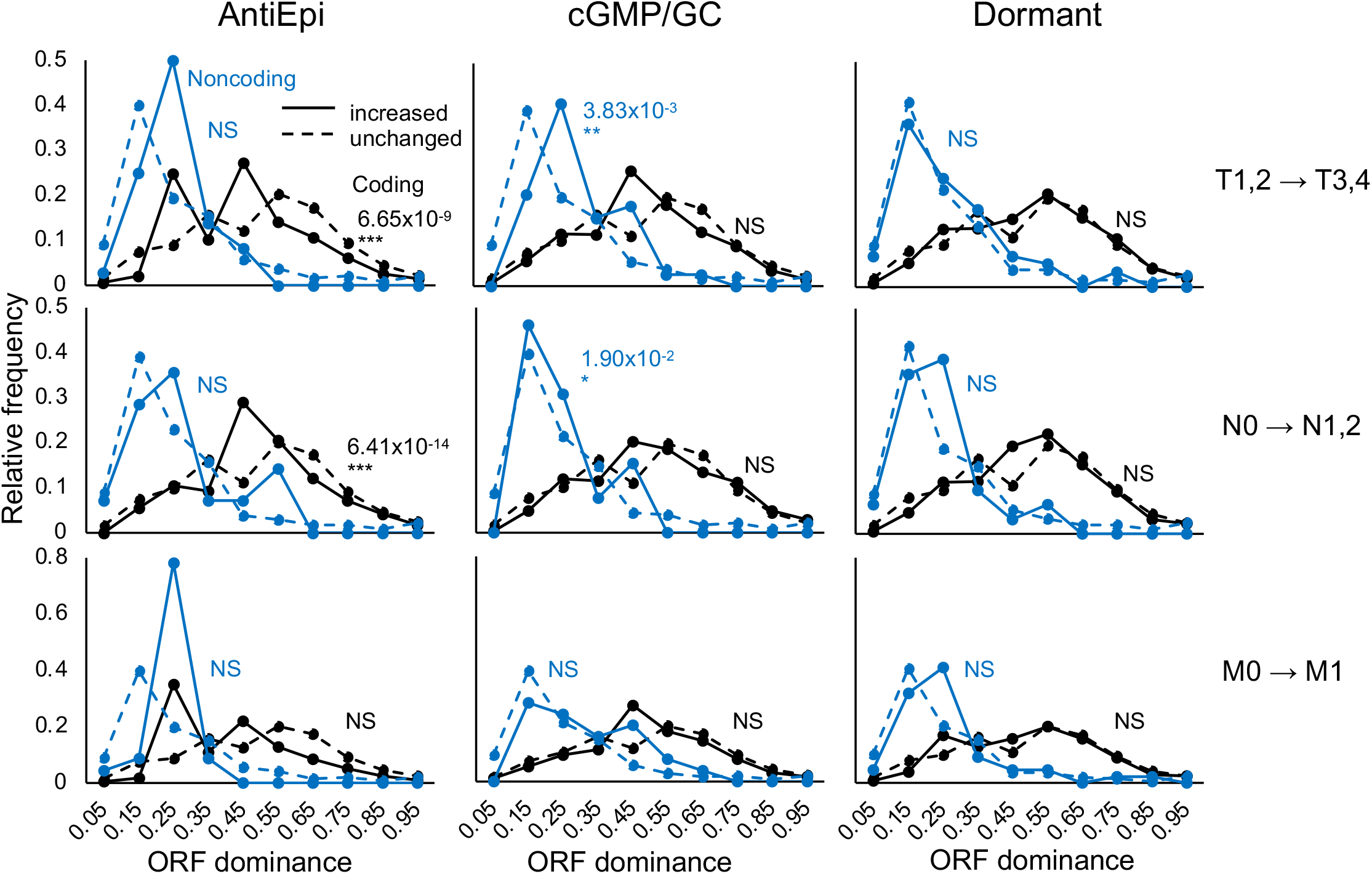
Relationship between ORF dominance distribution and TNM classification. ORF dominance distributions of transcripts with increased (solid line) or unchanged (dashed line) expression in late grades of T (top), N (middle), and M (bottom) in the three subgroups of human colorectal cancers. *P*-value was determined using the Mann–Whitney *U* test (***: *P* < 0.001, **: *P* < 0.01, *: *P* < 0.05, NS: not significant).

To test this hypothesis, we analyzed transcriptome data from a mouse skin carcinogenesis model (Aoto et al., 2017) that enables the observation of multi-step carcinogenesis from papilloma to metastatic cancer. In this model, skin cancer was induced by treating mouse skin with DMBA (7,12-dimethylbenz(a)anthracene) and TPA (12-*O*-tetradecanoylphorbol-13-acetone). Coding transcripts from normal skin (NORM), papilloma (PAPI), carcinoma (CARC), and metastatic tumors (META) obtained from the same mice were analyzed. We calculated the ORF dominance of transcripts with more than a two-fold upregulation in expression at the PAPI, CARC, and META stages (Figure S14, left). ORF dominance was significantly elevated during tumor initiation (NORM→PAPI), and the peak of the ORF dominance distribution shifted to lower values during tumor promotion (PAPI→CARC), while the distribution remained largely unchanged during tumor progression (CARC→META). The distribution of transcript length showed no significant changes (Figure S14, right).

## Discussion

In this study, we aimed to examine the role of ORF dominance in capturing the process of cancer evolution. In a previous report by Suenaga et al. (2022), we explored the association between ORF dominance calculated from transcriptome data of extant species and evolutionary phylogenetic trees. In this study, we compared ORF dominance between normal and cancerous tissues using publicly available patient-derived transcriptome data. We also investigated changes in ORF dominance during carcinogenesis and cancer progression using multiple mouse carcinogenesis models.

Our findings revealed an overall increasing trend in ORF dominance during carcinogenesis for both coding and noncoding transcripts (Figure 8). Gene ontology analysis of genes with elevated ORF dominance identified associations with the Golgi apparatus, endoplasmic reticulum, cell cycle, and mitosis. Conversely, decreased ORF dominance was associated with the mitochondrion, DNA damage, and DNA repair. Given that ORF dominance correlates with the translation frequency of ribosome-bound RNA, these fluctuations in gene translation likely contribute to carcinogenesis (Figure 8A). We observed high ORF dominance in noncoding RNAs that translate neoantigens, particularly with a peak around 0.25. Additionally, noncoding transcripts with increased expression in advanced human colorectal cancers exhibited a peak at 0.25 in ORF dominance. Therefore, noncoding RNAs with elevated ORF dominance could serve as promising candidates for RNA-based neoantigen translation and cancer immunotherapy in advanced cancers. Interestingly, elevations in ORF dominance were detected earlier in pancreatic organoids than in bile duct organoids. In mouse models of colorectal cancer and human colorectal cancers, elevated ORF dominance was associated with subgroups linked to metastasis (cGMP/GC and Dormant). These results suggest that increased ORF dominance may occur earlier in more aggressive cancer types or subtypes.

**Figure 8.**
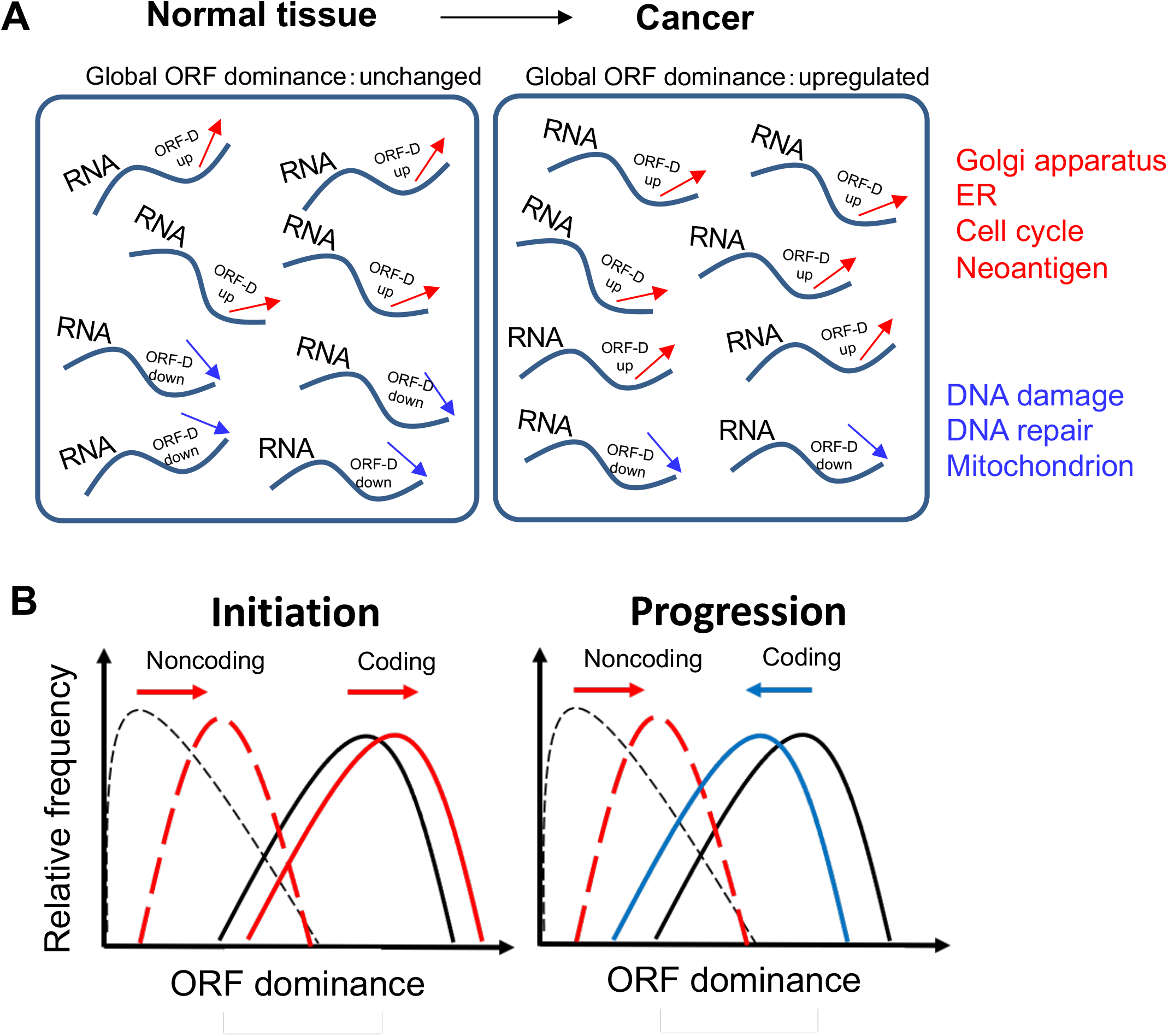
Model illustrating the relationship between ORF dominance and cancer initiation and progression. (A) Global changes in ORF dominance (ORF-D) during carcinogenesis and associated gene functions. (B) Changes in ORF dominance during cancer initiation and progression.

Regarding overall transcriptome changes, approximately 50% of transcripts exhibited altered ORF dominance during carcinogenesis. This finding aligns with previous reports on transcript alterations such as alternative splicing and alternative polyadenylation, which are prevalent in cancer (Climente-González et al., 2017; Vitting-Seerup et al., 2017). Abnormalities in genome-wide alternative polyadenylation in cancer have been reported to result in global 3’UTR shortening (Mayr et al., 2009; Mitschka et al., 2022; Desterro et al., 2020). The observed increase in ORF dominance in our mouse experiments, primarily driven by the shortening of sORF lengths, is consistent with the consequences of 3’UTR shortening. Moreover, the inactivation of DNA repair genes through intron polyadenylation (Dubbury et al., 2018) is in line with our findings that reduced ORF dominance was associated with DNA damage and DNA repair, primarily due to pORF shortening. Therefore, ORF dominance serves as a useful indicator of RNA isoform changes during cancer evolution and provides insights into their impact on translation.

Secondly, the elevation of ORF dominance during the oncogenic process was observed regardless of the presence of genetic mutations. Previous studies have attributed cancer-specific alternative splicing to dysregulated transcription rather than somatic mutations (Huang KK et al., 2021), and genetic mutations alone cannot explain splicing abnormalities (Kahraman et al., 2020), which aligns with our analysis using ORF dominance. Additionally, a previous report by Ono et al. (2021) suggested a decrease in the diversity of genomic mutations during Tp2 and Tp3 in a mouse colorectal cancer model, while the diversity of the transcriptome increased during the same time points. Although genetic mutations are major contributors to carcinogenesis, their contribution to changes in ORF dominance was not evident in our study.

Lastly, contrasting carcinogenesis, a decrease in ORF dominance of coding RNAs was observed during cancer progression in a skin carcinogenesis model and in a human colorectal cancer subgroup (AntiEpi). Conversely, an increasing trend was detected for noncoding RNAs during cancer progression in human colorectal cancers. These changes occurred prior to distant metastasis. In other words, coding and noncoding RNAs exhibited diminishing molecular distinctions during cancer progression before the development of metastatic tumors, blurring the boundary between them (Figure 8B). In our analysis of biological evolution, we proposed that the blurring of the coding noncoding RNA boundary is associated with a decrease in effective population size and an increased risk of extinction, while also providing opportunities for the emergence of new genes that enable species to adapt to new environments (Suenaga et al., 2022). The observation that the coding noncoding RNA boundary is also blurred during cancer progression suggests that the number of proliferative cancer cells (effective population size) in cancer tissue may decrease as cancer progresses before the development of metastatic tumors. This notion aligns with the fact that genes associated with the cell cycle and mitosis exhibited decreased ORF dominance in pancreatic cancer models (SC), and the Dormant subtype, characterized by low proliferative ability (low effective population size), demonstrated the highest metastatic potential with no significant changes in ORF dominance following its emergence.

In summary, our study elucidated the significance of changes in ORF dominance during cancer evolution as follows:

(1) ORF dominance undergoes extensive changes during carcinogenesis, influencing the translation efficiency and resulting in increased expression of oncogenes related to cell proliferation and decreased expression of genes involved in DNA damage response.

(2) Changes in ORF dominance during the carcinogenic process are not primarily driven by genetic mutations, but rather by other mechanisms such as epigenetic and/or transcriptional alterations.

(3) The boundary between coding and noncoding RNA becomes blurred prior to the development of metastatic tumors.

However, our study did not demonstrate whether genes affected by changes in ORF dominance actually play a role in carcinogenesis and cancer progression, which is a question that needs to be addressed in future research. The upstream regulatory mechanisms governing alterations in ORF dominance remain unknown. Therefore, accumulating and analyzing experimental data using ORF dominance as a starting point will be crucial for a deeper understanding of carcinogenesis and cancer progression.

## Acknowledgments

We gratefully acknowledge the valuable comments from Toshinori Ozaki, as well as the technical assistance provided by Kyoko Takahashi, Noriko Miyazaki, Akiko Endo, and Aya Washio. We also thank Elaine Fuchs for providing LV-Cre pLKO.1. This work received partial support from the Takeda Science Foundation (Y.S.) and e-ASIA Grant 21jm0210092h0001 from the Japan Agency for Medical Research and Development (Y.H.).

The results presented in this study are based in part on data generated by the TCGA Research Network: https://www.cancer.gov/tcga.

## Author contributions

Y.S. and H.K. conceived and designed the research plan. Y.S., H.K., K.H., J.L., K.K., Y.K., E.F., K.O., E.K., Y.W., M.Kat., M.Kaw., and Y.H. performed data analysis. Y.S., H.K., J.L., Y.H., K.K., Y.K., M.Kat., and Y.H. contributed to writing the manuscript.

## Additional information

Supplementary information is available.

## Data availability

List of sequencing sources can be found in Table S17. Source data for analyses and figures can be accessed from the DNA Data Bank of Japan (DDBJ; www.ddbj.nig.ac.jp) with accession No. PRJDB15934.

