## Supplemental Figures for "Global changes in open reading frame dominance of RNAs during cancer initiation and progression"

$$\text{ORF dominance} = \frac{\text{Length of primary ORF}}{\text{Sum of all ORF lengths}}$$

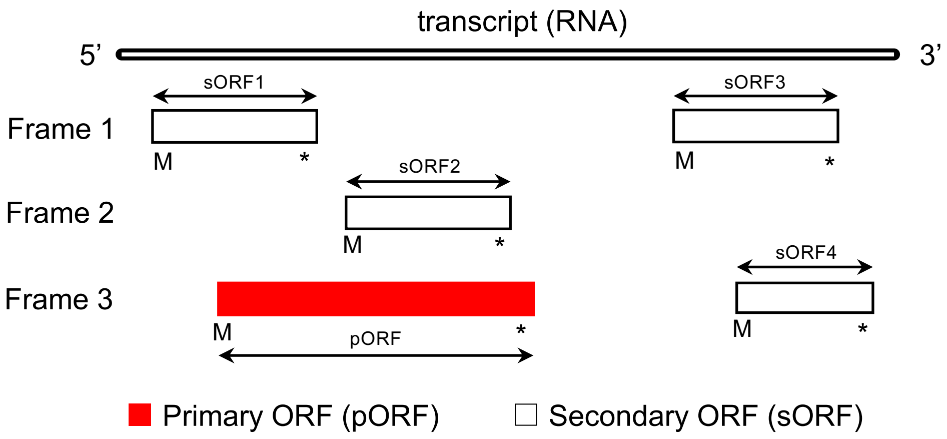


**Figure S1. Definition of ORF dominance.** Definition of ORF dominance is presented in the upper panel, while the lower panel provides a conceptual schematic representation of ORFs in the three reading frames of an RNA. The primary ORF, indicated by red rectangles, is the longest ORF, while the secondary ORFs, represented by white rectangles, include all other ORFs. ORF length is defined as the length of the amino acid sequence, excluding the stop codon (*). “M” indicates the start codon.

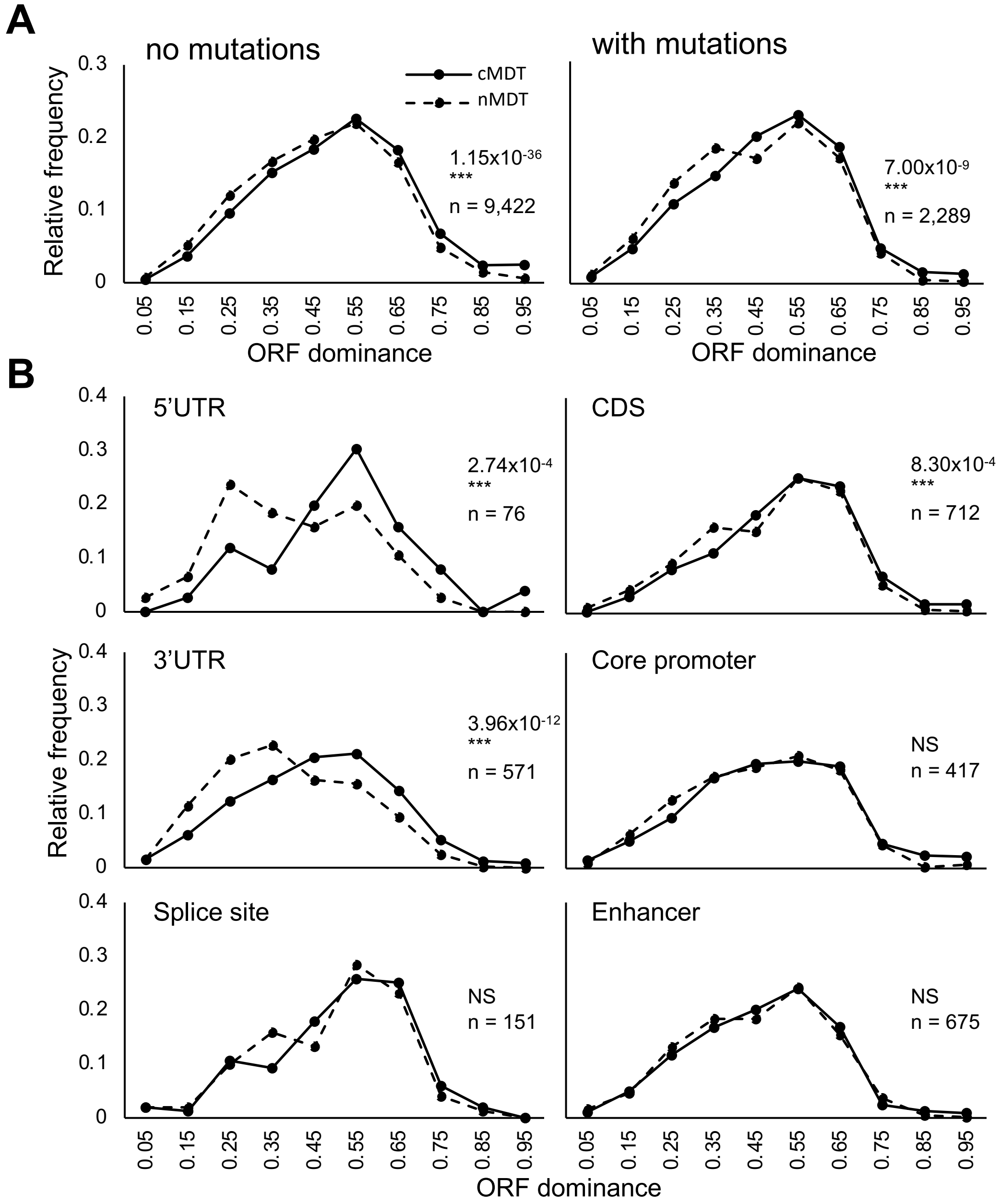


**Figure S2. Relationship between ORF dominance and somatic mutations.** (A) Distribution of ORF dominance in cMDT (solid line) and nMDT (dashed line) with or without mutations. (B) ORF dominance distribution of cMDT mutated in different genomic regions. *P*-value was calculated using the Mann-Whitney *U* test (***: $P < 0.001$, **: $P < 0.01$, *: $P < 0.05$, NS: not significant).

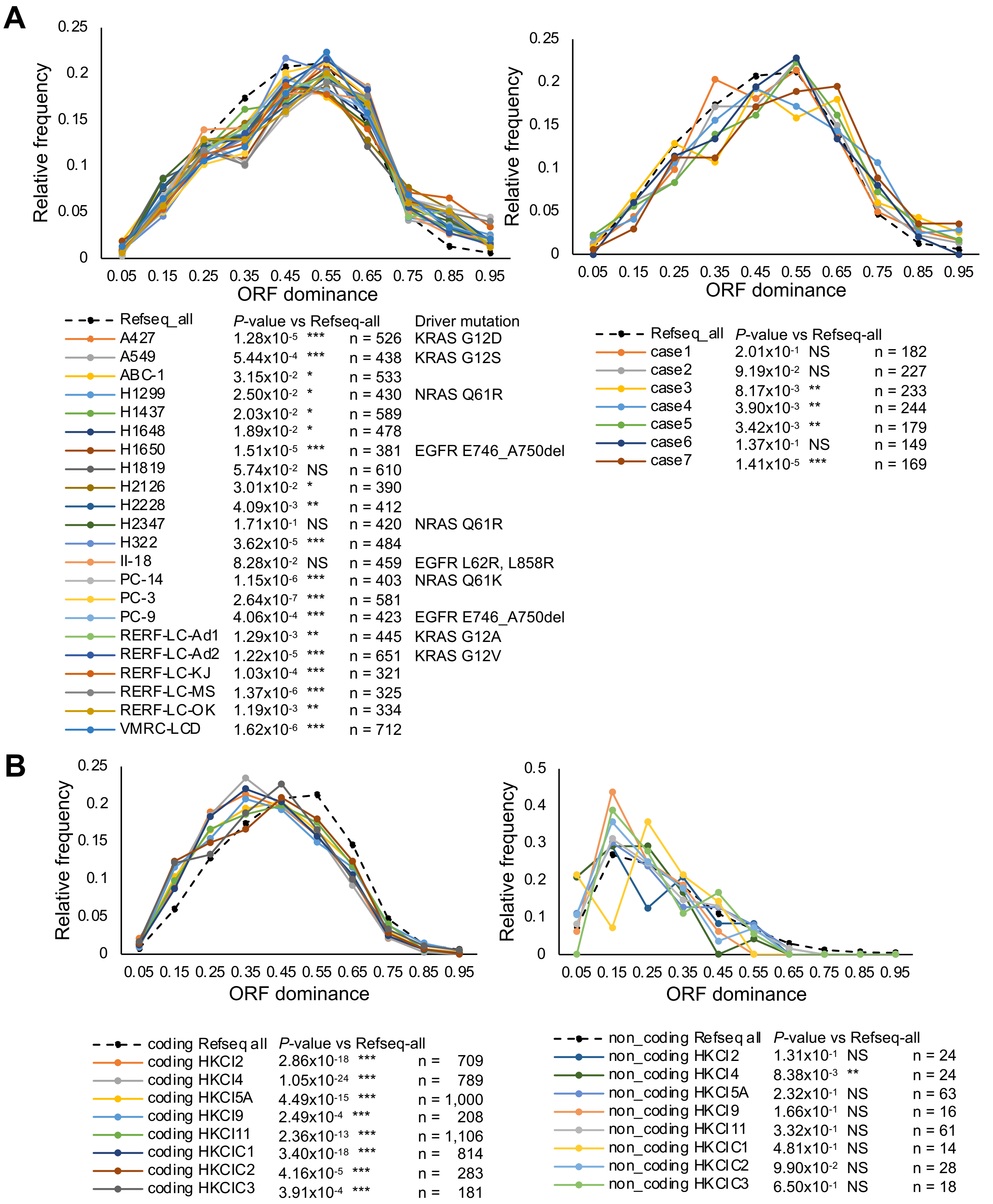


**Figure S3. LR-seq-based ORF dominance distribution of lung and liver cancer.** (A) ORF dominance distribution of non-small cell lung cancer-derived cell lines (left) and patient specimens (right). (B) ORF dominance distribution of hepatocellular carcinoma patient specimens, coding (left) and noncoding (right). *P*-value was calculated using the Mann-Whitney *U* test (***: $P < 0.001$, **: $P < 0.01$, *: $P < 0.05$, NS: not significant).

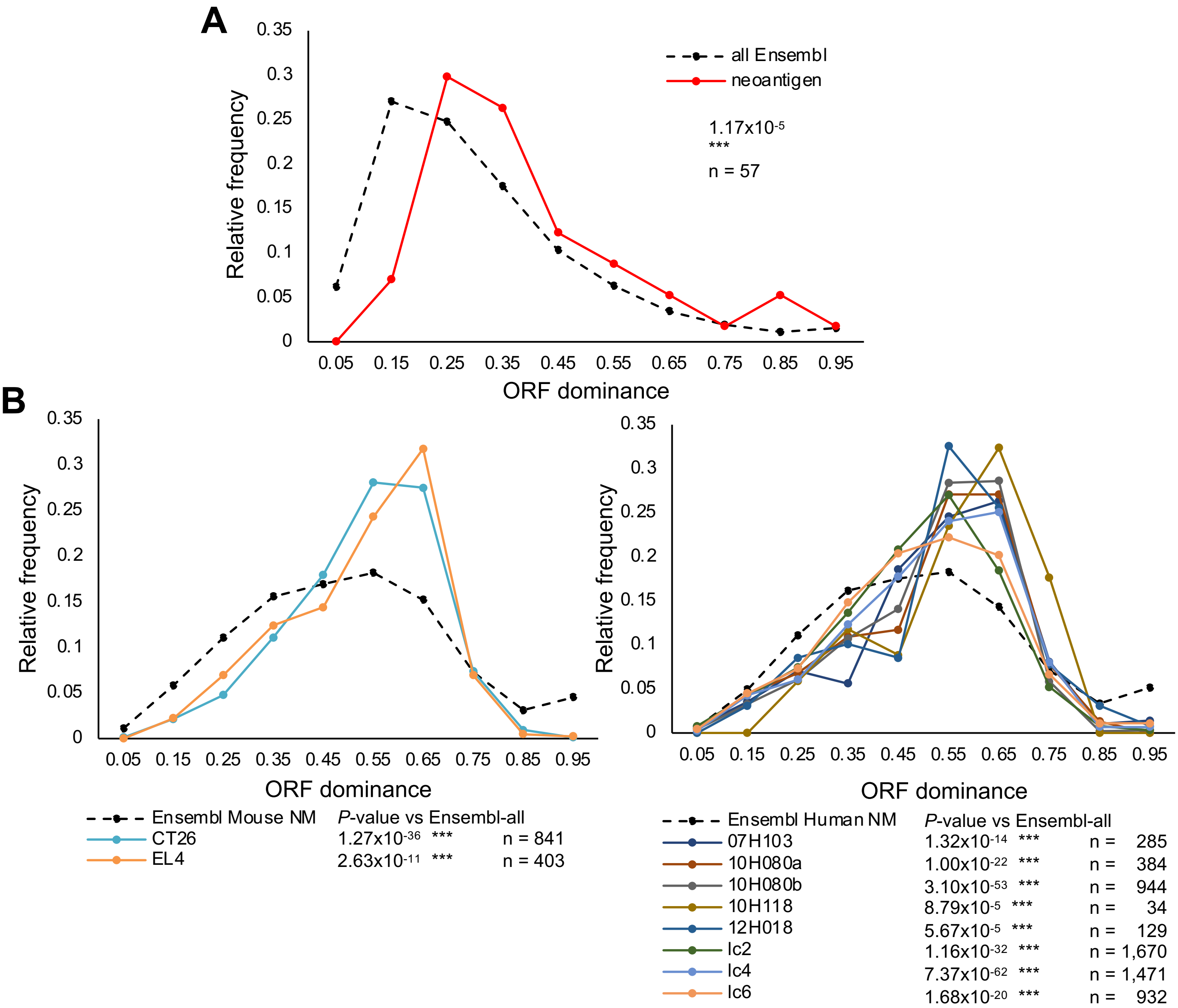


**Figure S4. ORF dominance distribution of noncoding RNAs producing neoantigens.** (A) ORF dominance distribution of noncoding RNAs producing potential tumor neoantigens (red solid line). (B) ORF dominance distribution of tumor-specific antigens in murine cell lines (left) and human primary tumors (right). *P*-value was calculated using the Mann-Whitney *U* test (***: $P < 0.001$, **: $P < 0.01$, *: $P < 0.05$, NS: not significant).

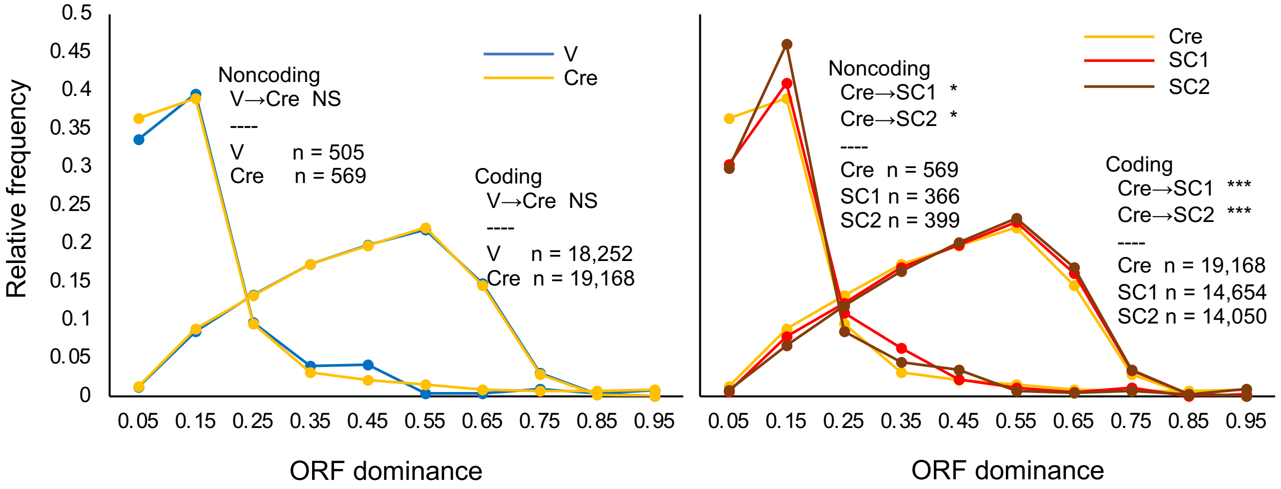


**Figure S5. ORF dominance distribution of bile duct organoids using the average value as a representative value for ORF dominance calculation.** V: vehicle, Cre: mutation-induced organoid by Cre recombinase. SC1/2: Subcutaneous tumor-derived organoid. *P*-value was determined using the Mann-Whitney *U* test (***: $P < 0.001$, **: $P < 0.01$, *: $P < 0.05$, NS: not significant).

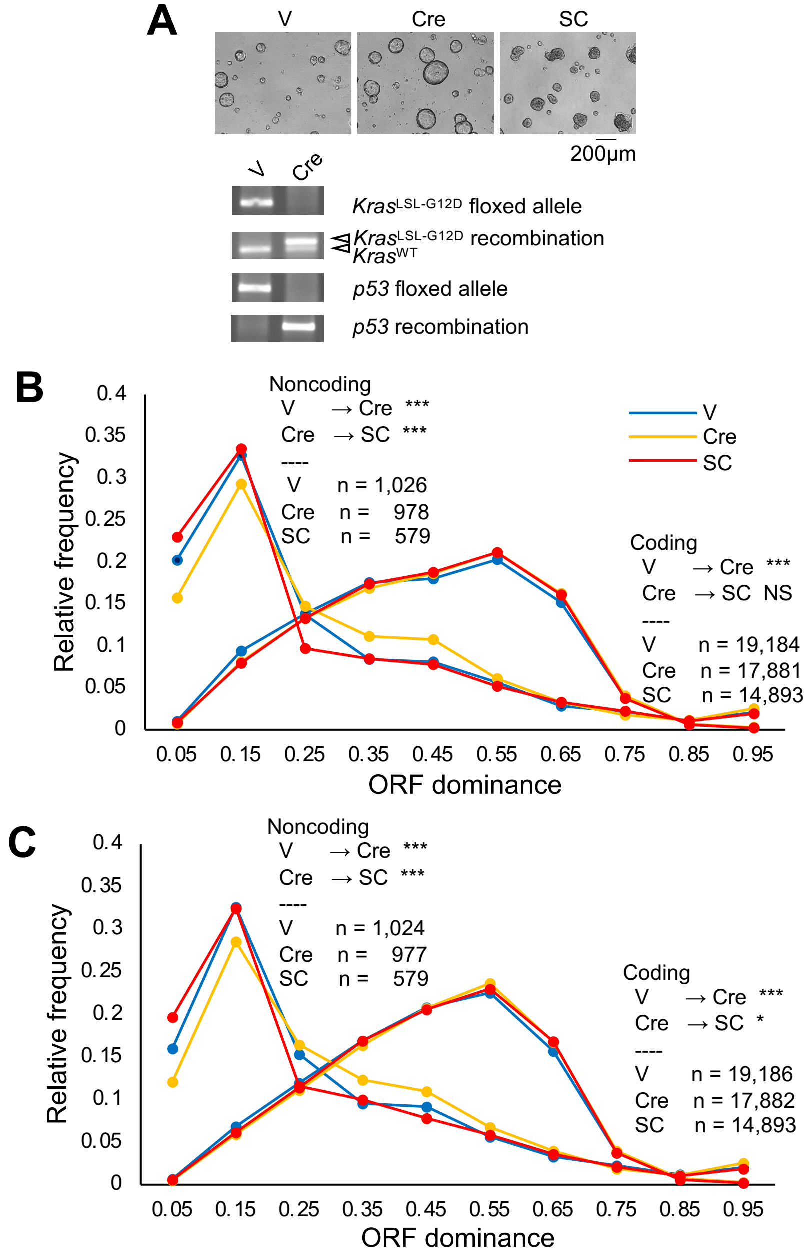


**Figure S6. ORF dominance distribution in a mouse pancreatic carcinogenesis model.** (A) Phase contrast images of mouse organoids and genetic PCR analysis focused on *Kras* and *p53* loci. ORF dominance distribution of transcripts based on LR-seq data using the longest transcript (B) or average value (C) for the calculation of ORF dominance. *P*-value was calculated using the Mann-Whitney *U* test (***: $P < 0.001$, **: $P < 0.01$, *: $P < 0.05$, NS: not significant).

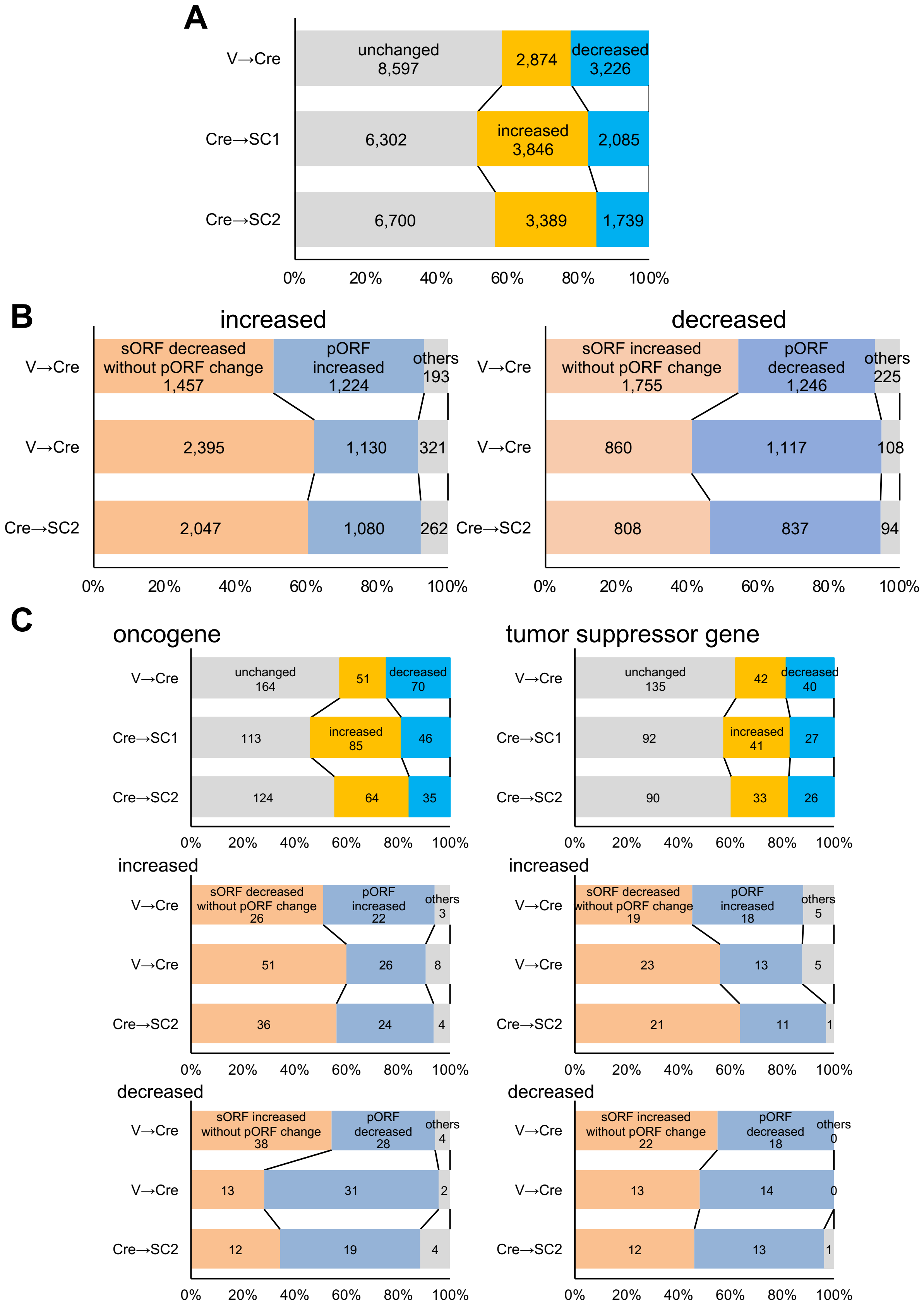


**Figure S7. Causes of ORF dominance changes in cancer organoids.** (A) Percentage of transcripts with increased, decreased and unchanged ORF dominance. The number of RNAs is indicated. (B) Percentage of the causes for increased and decreased ORF dominance. (C) Similar analysis for oncogenes (left) and tumor suppressor genes (right), respectively.

A

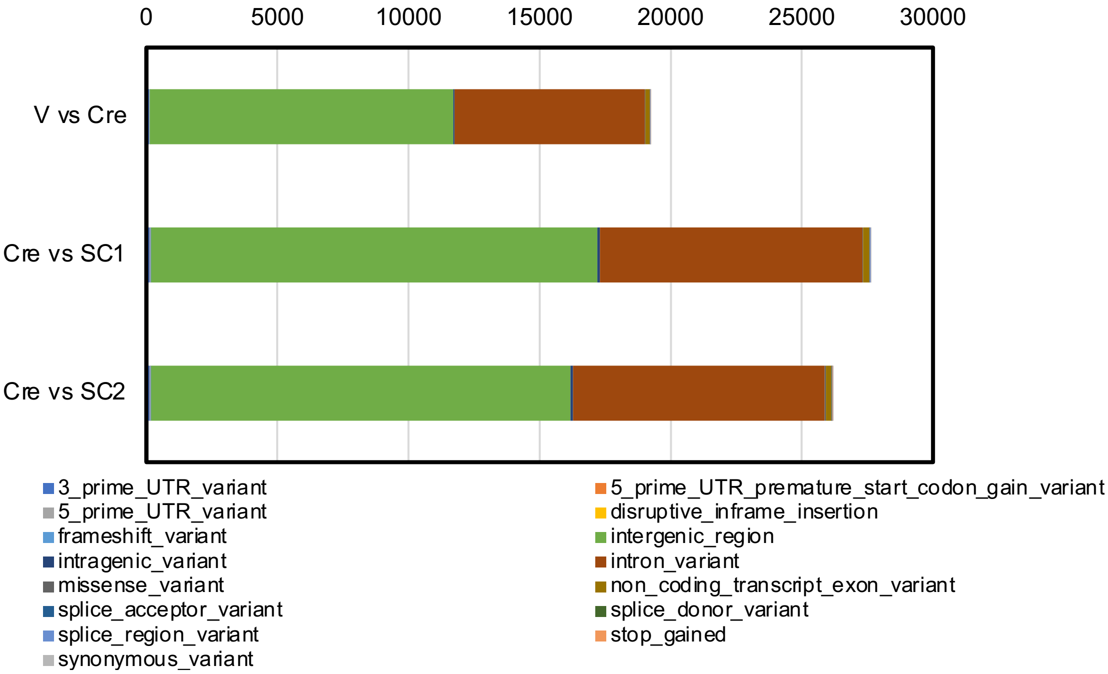


B

| Mutation type | V vs Cre | Cre vs SC1 | Cre vs SC2 |
|---|---|---|---|
| 3_prime_UTR_variant | 108 (0.56%) | 139 (0.50%) | 136 80.52%) |
| 5_prime_UTR_premature_start_codon_gain_variant | 0 (0.00%) | 1 (0.00%) | 1 (0.00%) |
| 5_prime_UTR_variant | 22 (0.11%) | 24 (0.09%) | 28 (0.11%) |
| disruptive_inframe_insertion | 0 (0.00%) | 0 (0.00%) | 0 (0.00%) |
| frameshift_variant | 3 (0.02%) | 3 (0.01%) | 5 (0.02%) |
| intergenic_region | 11,559 (60.12%) | 17,042 (61.66%) | 15,991 (61.06%) |
| intragenic_variant | 61 (0.32%) | 87 (0.31%) | 101 (0.39%) |
| intron_variant | 7,254 (37.73%) | 10,004 (36.20%) | 9,609 (36.69%) |
| missense_variant | 28 (0.15%) | 29 (0.10%) | 35 (0.13%) |
| non_coding_transcript_exon_variant | 167 (0.87%) | 248 (0.90%) | 233 (0.89%) |
| splice_acceptor_variant | 1 (0.01%) | 5 (0.02%) | 4 (0.02%) |
| splice_donor_variant | 0 (0.00%) | 3 (0.01%) | 2 (0.01%) |
| splice_region_variant | 14 (0.07%) | 29 (0.10%) | 27 (0.10%) |
| stop_gained | 2 (0.01%) | 3 (0.01%) | 2 (0.01%) |
| synonymous_variant | 9 (0.05%) | 20 (0.07%) | 17 (0.06%) |

**Figure S8. Somatic mutations detected in whole-genome sequencing of mouse bile duct organoids.** The number and composition of mutations detected by comparing genomic DNA sequences between the indicated bile duct organoids are shown in bars (A) and a table (B).

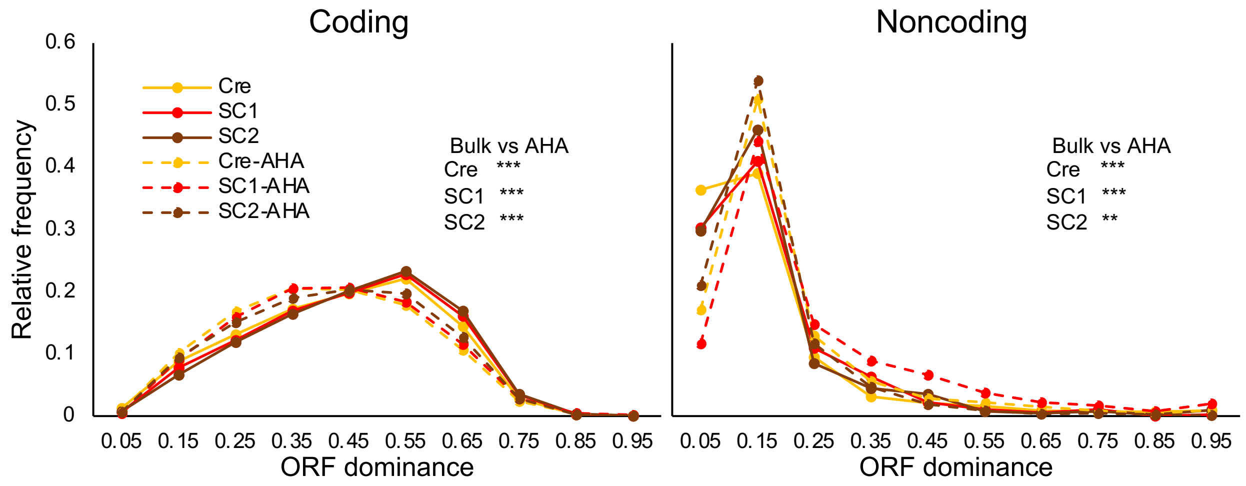


**Figure S9. ORF dominance distribution of transcripts bound to ribosomes in a mouse bile duct carcinogenesis model using the average value for the calculation of ORF dominance.** ORF dominance distributions of coding (left) and noncoding (right) transcripts bound to ribosome complexes (AHA) in mouse bile duct organoids. Bold and dotted lines indicate all RNAs detected in the organoids (bulk, same data as Figure 2D) and RNAs detected after AHARIBO (AHA), respectively. Statistical significance was determined using the Mann-Whitney *U* test (***: $P < 0.001$, **: $P < 0.01$, *: $P < 0.05$, NS: not significant).

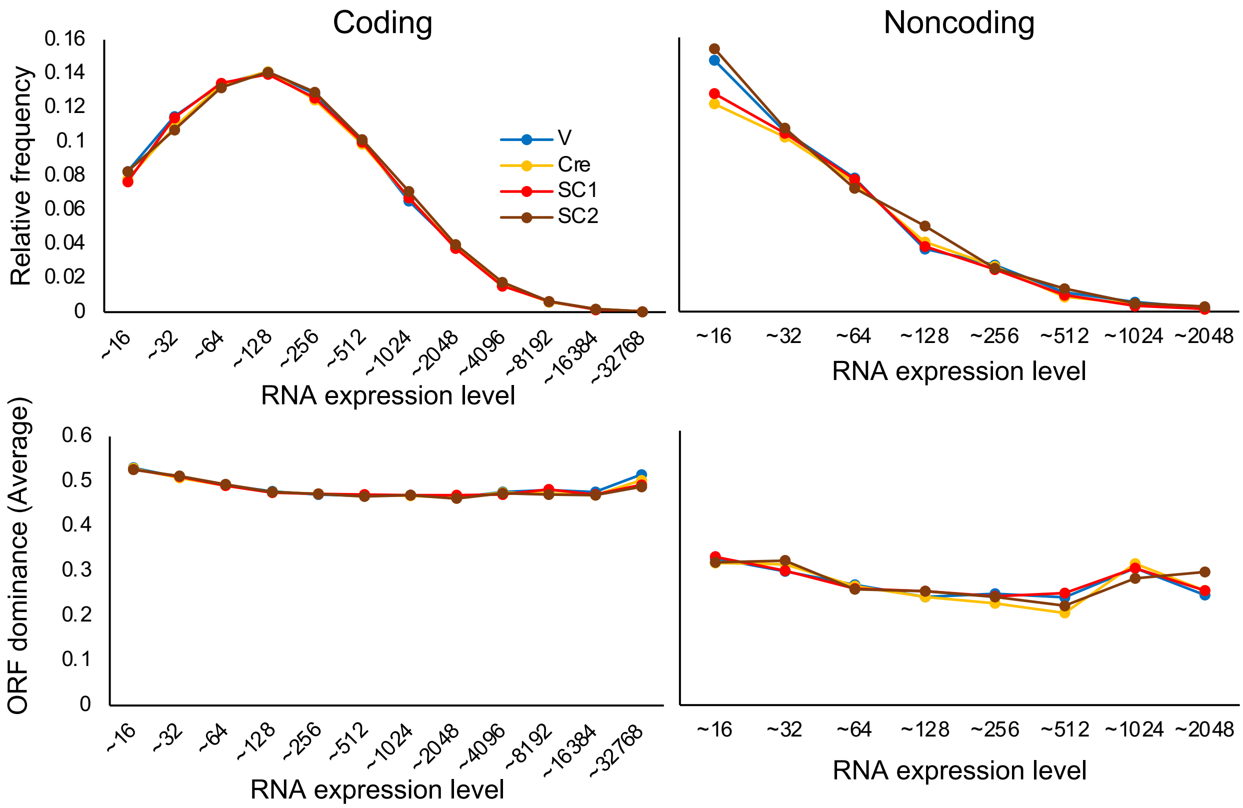


**Figure S10. Relationship between ORF dominance and RNA expression level.** Distribution of relative frequencies of RNA expression levels in coding (upper left) and noncoding (upper right) RNA. The relationship between average values of ORF dominance and RNA expression levels in coding (lower left) and noncoding (lower right) RNAs.

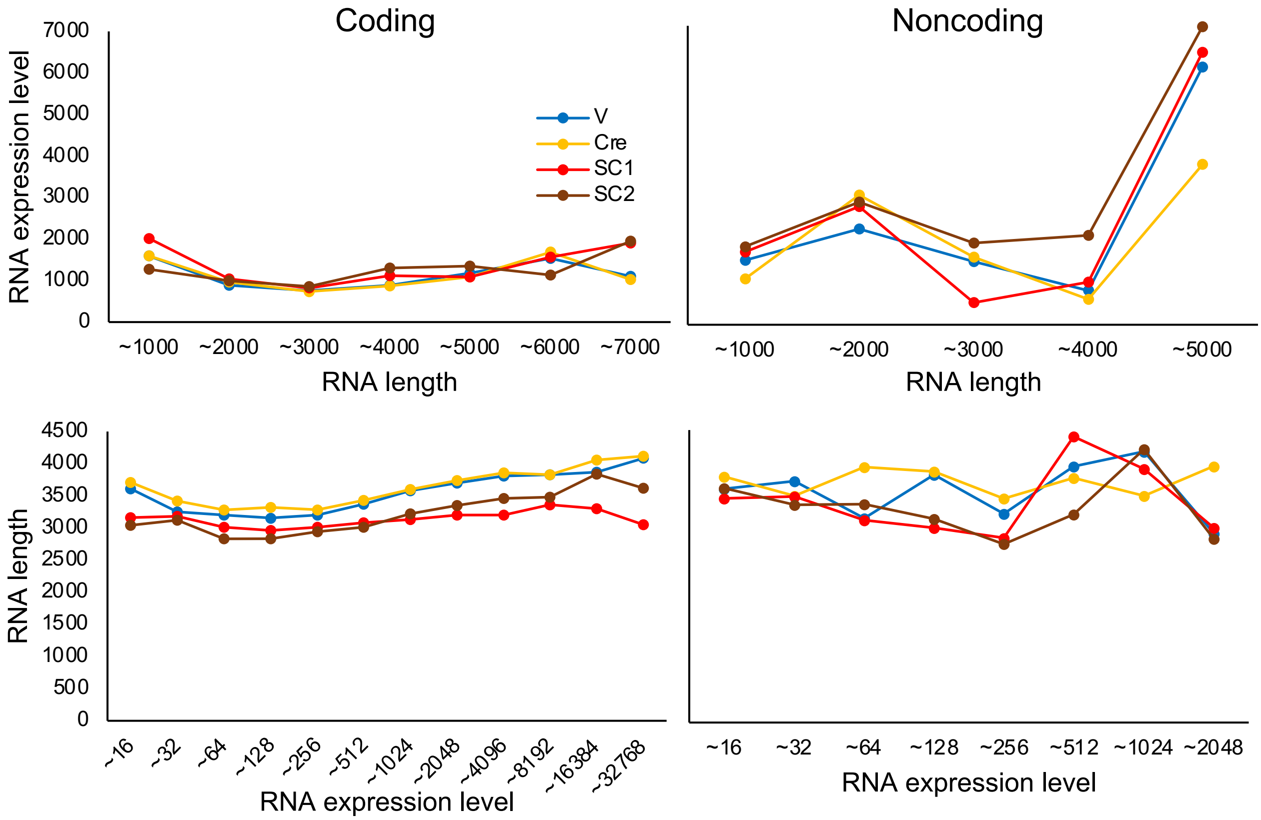


**Figure S11. Relationship between RNA length and RNA expression level in mouse bile duct organoids.** Average values of RNA expression levels and RNA length (nucleotides) were used in the upper and lower panels, respectively.

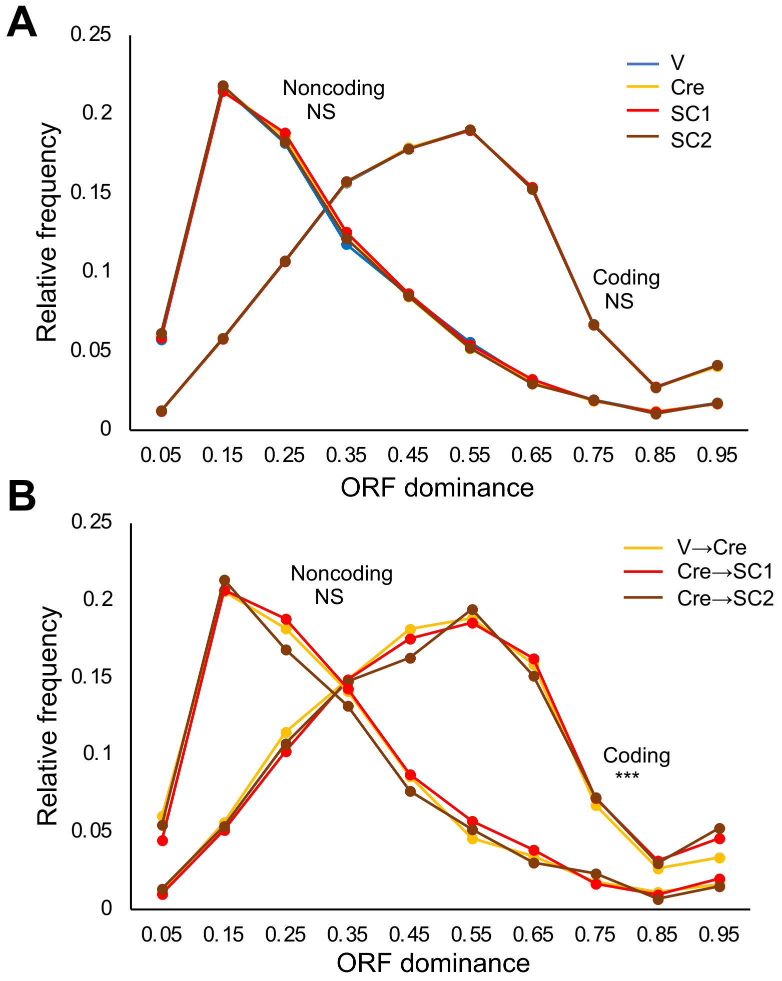


**Figure S12. ORF dominance distribution of transcripts expressed in mouse organoids based on SR-seq data.** (A) ORF dominance distribution using all transcript data. (B) ORF dominance distribution of transcripts with elevated expression. Statistical significance was determined using the Mann-Whitney *U* test (***: $P < 0.001$, NS: not significant).

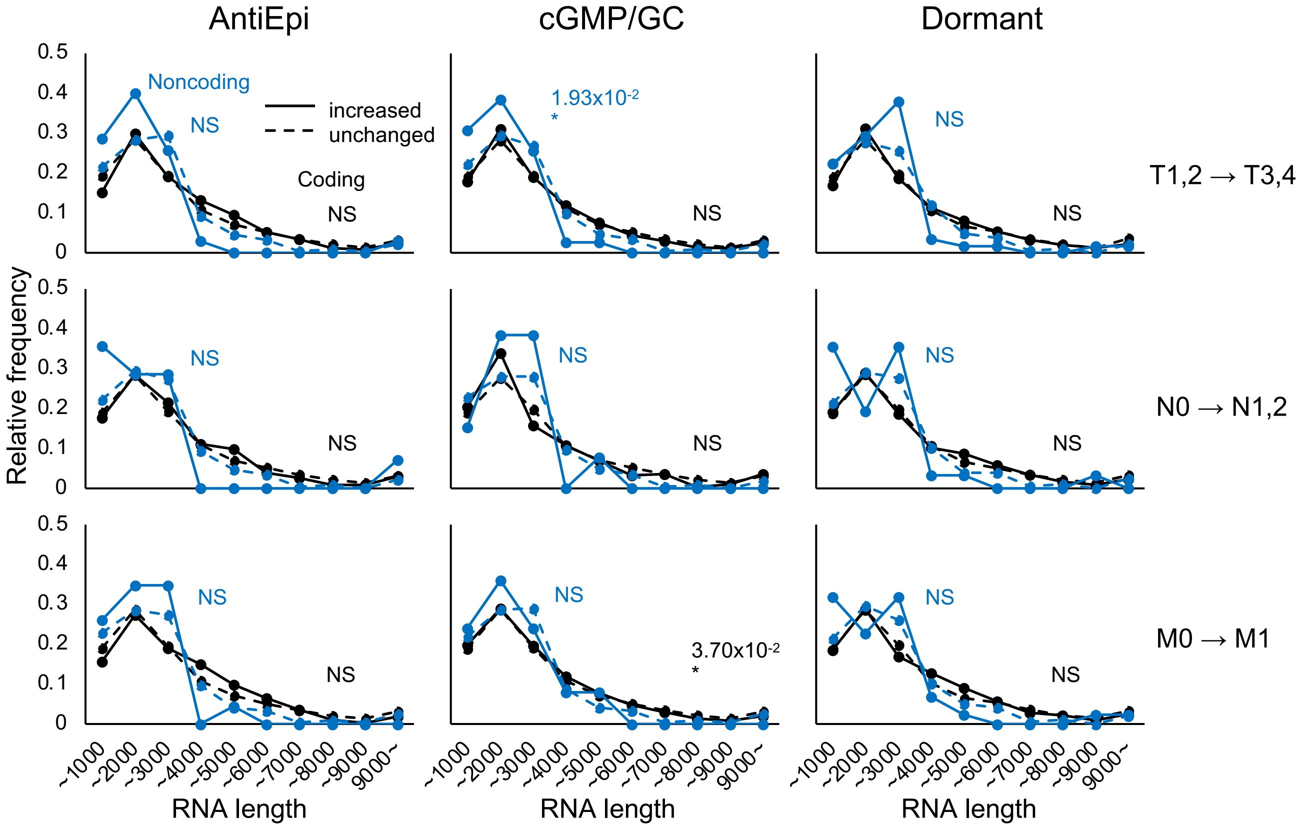


**Figure S13. Relationship between RNA length distribution and TNM classification.** Transcript length (nucleotide) distributions of transcripts with increased (solid line) or unchanged (dashed line) expression in late grades of T (top), N (middle), and M (bottom) in the three subgroups of human colorectal cancers. *P*-value was calculated using the Mann-Whitney *U* test (*: $P < 0.05$, NS: not significant).

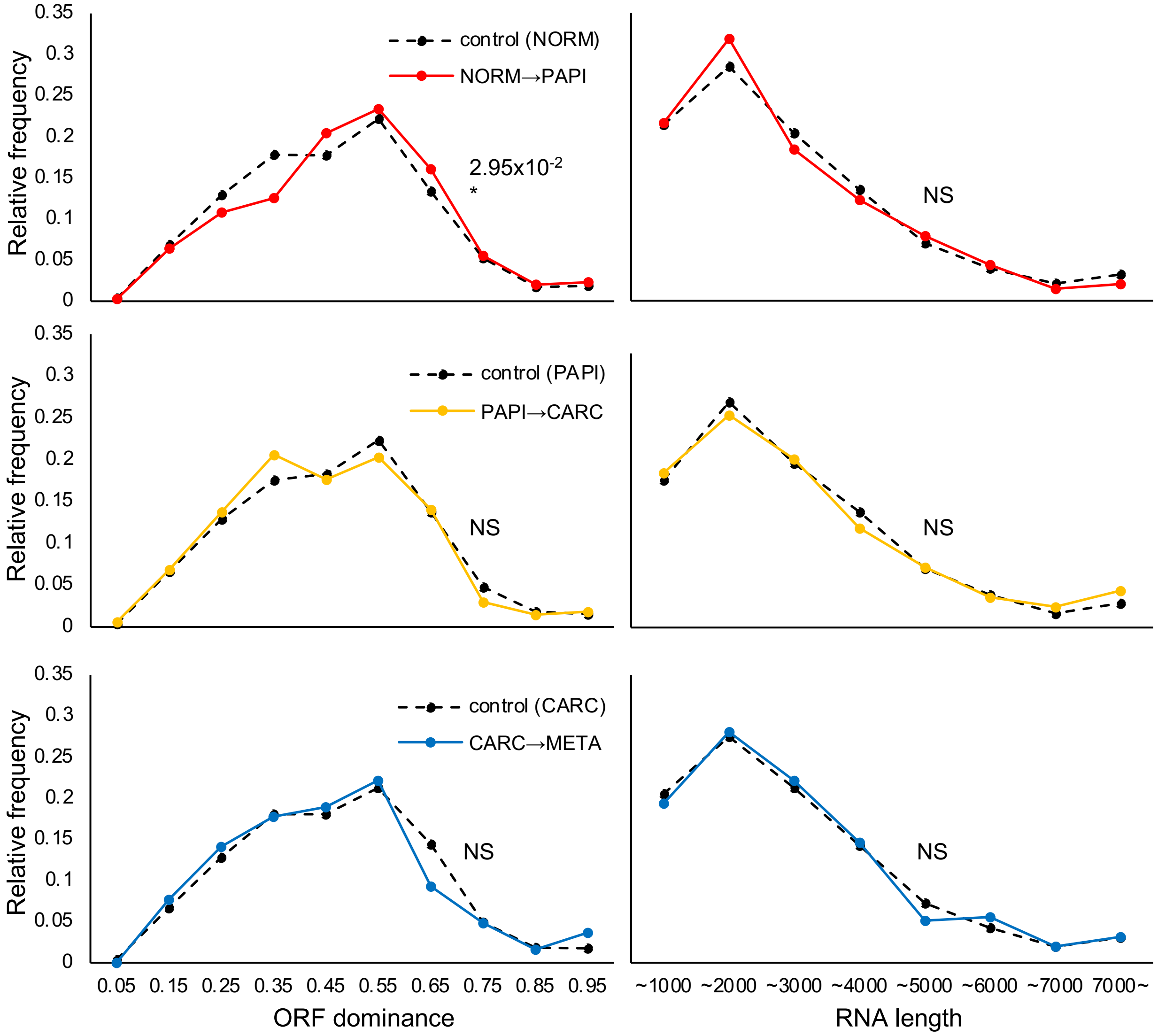


**Figure S14. ORF dominance distribution of transcripts with elevated expression during cancer evolution in a mouse skin carcinogenesis model.** ORF dominance (left) or RNA length (right) distributions of transcripts with elevated expression levels in later stages (solid lines) were compared with those of transcripts detected in earlier stages (dashed line). *P*-value was calculated using the Mann-Whitney *U* test (*: $P < 0.05$, NS: not significant).
